# Root-pore interactions, the underestimated driver for rhizosphere structure and rhizosheath development

**DOI:** 10.1101/2024.12.06.627175

**Authors:** Maik Geers-Lucas, Andrey Guber, Alexandra Kravchenko

## Abstract

Physical characteristics of rhizosphere and rhizosheath, i.e. root-adhering soil, are crucial for plant performance. Yet, the drivers of the rhizosphere’s structural properties and their relationships with rhizosheath development remain unclear.

We used X-ray computed micro-tomography (i) to explore two drivers of rhizosphere porosity: root-induced changes vs. preferential root growth into soil with certain pore characteristics and (ii) to estimate their contributions to rhizosphere macroporosity gradients and rhizosheath formation. Rhizosheath development was assessed in relation to rhizosphere macroporosity and rhizodeposition after ¹⁴C labeling.

Our results confirmed that both root-induced changes and growth preferences shape rhizosphere structure, with their relative significance depending on the inherent macropore availability. In intact soils, growth preferences were the dominant factor, while in sieved soils the root-induced changes became equally important. Rhizosheath formation was associated with roots compacting their surrounding and releasing carbon. However, no correlation was found between rhizosheath formation and the actual rhizosphere, i.e., the volume of soil adjacent to the roots.

The study offers new process-level understanding of rhizosphere porosity gradients, while emphasizing caution in interpreting root growth data from sieved soil studies. Similarly, traditional destructively sampled rhizosheath may not fully capture the true characteristics of the actual rhizosphere, underscoring importance of intact-soil analyses.

## 1 Introduction

Rhizosphere, the narrow zone of soil directly surrounding plant roots, is one of the most dynamic regions within the soil that frequently undergoes rapid and significant alterations in its physical, chemical, and biological properties (Hinsinger *et al*., 2009). Modifications in soil structure, i.e. arrangement of soil solids and pores, are arguably the most prominent and impactful of such root-induced changes. The rhizosphere is the bottleneck through which water and nutrients must flow when they are either released or taken up by the plant root, and these fluxes occur via soil pores. Thus, structural changes in characteristics of pore and solid components forming the rhizosphere have a direct impact on water flow, nutrient cycling, and carbon storage (Aravena *et al*., 2011; van Veelen *et al*., 2019; Landl *et al*., 2021; Pankievicz *et al*., 2022; Lucas *et al*., 2023b).

The rhizosphere pore structure is often reported to differ significantly from that of bulk soil (Helliwell *et al*., 2019; Koebernick *et al*., 2019; Lucas *et al*., 2019), i.e., the soil not influenced by the roots. However, the findings are often contradictory. For example, several works that used X-ray computed tomography (X-ray µCT), currently the most advanced approach for *in situ* visualization of plant roots and their contact with the soil, reported reduction in soil macroporosity within the rhizosphere (Feeney *et al*., 2006; Helliwell *et al*., 2017, 2019; Rabbi *et al*., 2018a,b; Zhang *et al*., 2020), while many others noted an increase (Bruand *et al*., 1996; Vollsnes *et al*., 2010; Lucas *et al*., 2019). Such discrepancies are indicative of the current lack of knowledge of the underlaying mechanisms, which substantially limit the progress in understanding and modeling plant-soil interactions (van Veelen *et al*., 2019; Landl *et al*., 2021; Schnepf *et al*., 2022). There is a pressing need for a conceptual and quantitative framework to guide future experimental designs and the interpretation of root-soil interaction data. In this study, we aim to lay the experimental groundwork for developing such a framework.

We surmise that the reported contradictions in porosity gradients around the roots result from simultaneous contributions of two disparate mechanisms: 1) direct root-induced changes and 2) indirect effects of root growth preferences. Direct changes take place when roots compress the soil while moving into and growing through relatively small pores, hence reducing the microporosity in the newly formed rhizosphere (Dexter, 1987). Later, the initial reduction can be somewhat abated due to structural changes generated by combined impacts of rhizodeposition, enhanced wetting/drying cycles in vicinity of the roots (Koebernick *et al*., 2018), root shrinkage during soil drying, and the “surface-wall effect” (Carminati *et al*., 2013; Koebernick *et al*., 2019). The latter refers to the packing of round-shaped soil particles against the growing root, which can increase macroporosity at the root surface (Koebernick *et al*., 2019).

The indirect outcomes arise from roots often preferring to grow into areas of low penetration resistance, such as large macropores (White & Kirkegaard, 2010; Lucas *et al*., 2019, 2022). In such cases, root growth might have a minimal impact on the surrounding soil structure. However, the macropores themselves often already contain greater proportions of smaller pores in their vicinity (Lucas *et al*., 2019). Roots populating such macropores will have a highly porous rhizosphere, due to existing soil pore structure and root growth preferences within it, rather than being an outcome of active soil modifications by the roots.

We argue that root growth preferences play a more critical role in shaping the rhizosphere’s pore structure than direct effects such as compressing of soil by the roots, making the initial pore structure the primary factor influencing the rhizosphere’s physical properties. Notably, all of the past experiments focusing on rhizosphere pore structure were conducted in artificially structured, e.g., sieved or mixed, soils. Such soil mixtures may not fully capture the complex interactions between roots and soil structure in natural, i.e., intact soil, environments. The intact soil is often abounded with large cracks of non-biological origin, biopores, and channels previously created by roots or living organisms (Lucas *et al*., 2023b; Vogel *et al*., 2024), which make it a substantial experimental challenge to identify new roots and quantify their rhizosphere macroporosity. Since both root-induced changes and root growth preferences may lead to similar variations in rhizosphere macroporosity, it is challenging to separate the contributions of these two processes. Distinguishing between them, however, is crucial for modeling and predicting the effects of roots on soil pore structure and for developing process-based approaches to improving rhizosphere properties.

The structure of the rhizosphere soil is also represented by its solid components, e.g., by sizes and volumes of soil particles within it. That is often investigated in a form of a rhizosheath, which is currently operationally defined as all soil adhering to the roots and which is considered as a practical way to sample and analyze the rhizosphere soil (Hallett *et al*., 2022). Rhizosheath formation and soil aggregation within it are influenced by a variety of factors, including root mucilage, root hairs, and wetting-drying cycles (Aslam *et al*., 2022; Cheraghi *et al*., 2023; Mo *et al*., 2023). Originally, the term rhizosheath was introduced for desert soils and referred to a distinct, sheath-like soil layer, created by rhizodeposits gluing soil particles together during drying and bound to the root surface (Price, 1911; Watt *et al*., 1994; Pang *et al*., 2017; Ndour *et al*., 2020; Mo *et al*., 2023). As part of the rhizosphere, rhizosheath soil is critical for nutrient acquisition and water retention, serving as a protective layer that increases root surface area suitable for performing these functions (Hallett *et al*., 2022; Cheraghi *et al*., 2023; Mo *et al*., 2023; Mueller *et al*., 2024). Rhizosheaths are therefore thought to buffer plants against abiotic stress, such as drought, by preventing air gaps around roots and maintaining optimal soil moisture levels (Hallett et al., 2022). While root-adhering soil constitutes just a fraction of the entire rhizosphere, it is commonly regarded as representative of the rhizosphere (Pang *et al*., 2017). However, despite extensive research on rhizosheath formation, the relationship between the rhizosheath and the actual rhizosphere remains unclear, leaving the question of whether root-attached soil accurately reflects the broader rhizosphere properties unanswered.

This study has two interconnected goals: (i) to quantify the contributions of the two mechanisms defining rhizosphere pore structure, i.e., root-induced changes and inherent root growth preferences; and (ii) to investigate relationships between these two mechanisms and rhizosheath development, specifically focusing on the role of heterogeneity in soil pore structure and plant growth preferences on its formation. To achieve these goals, we made use of previously published data (Lucas *et al*., 2023b), which involved Switchgrass (*Panicum virgatum* L.) and Black-Eyed-Susan (*Rudbeckia hirta* L.) grown in four types of ingrowth cores: intact and sieved cores from switchgrass and prairie systems. We differentiated the mechanisms defining rhizosphere macropore structure by analyzing images from repeated X-ray µCT imaging (before and after plant growth) and linked rhizosheath measurements via conventional soil sampling with the rhizosphere volume determined via X-ray µCT and assumed to represent the actual rhizosphere in the intact soil settings.

We hypothesize, first, that in intact soils, the contribution of root-induced changes to formation of rhizosphere pore structure will be minimal as roots will primarily reuse existing macropores, with macroporosity gradients largely driven by root growth preferences. Second, increased root growth into the soil matrix, accompanied by densification of rhizosphere structure and greater amounts of root-derived carbon inputs, will enhance soil adhesion to the roots, thereby promoting rhizosheath formation. Third, the size of the rhizosheath is representative of the size of the actual rhizosphere, which will be reflected in positive associations between the rhizosphere volume determined via X-ray µCT and the rhizosheath mass.

## 2 Methods

### 2.1 Experimental design and ^14^C labeling

This study analyzed images from an experiment reported by Lucas et al. (2023b), which provides detailed information about the experimental settings and plant growth conditions. Briefly, a split pot experiment was conducted using ingrowth cores (5 cm diameter) with different soil origins and pore structures (Fig. 1). The cores were collected from the DOE-Great Lakes Bioenergy Research Center’s (GLBRC) Marginal Land Experiment (MLE) site, established in 2013 in Oregon, Wisconsin. The plots sampled for this study had a long-term (7 years) history of disparate vegetation, namely, monoculture switchgrass (*Panicum virgatum* variety Cave-in-Rock) and restored prairie (an 18-species mix), the plant systems known for their roles in developing contrasting pore structures (Kravchenko *et al*., 2019; Li *et al*., 2024). A total of 60 intact cores were taken from the replicated experimental plots of both plant systems, with half of the cores kept intact and the rest sieved and repacked to the same bulk density. To ensure that the intact and sieved cores differed from each other only in terms of pore structure, the sieved soil was packed to the same bulk density as the intact cores and all >2 mm inclusions, such as stones and large root residues were incorporated back into the sieved soil prior to packing. We then placed four ingrowth cores into each of twelve pots (15 cm diameter, 20 cm height), with six replicated pots per plant species. Hoagland-Solution (Hoagland & Arnon, 1950) was added (1L per pot) while filling the pots with the soil as a basal fertilizer and the volumetric soil water content was set to 30% (corresponding to −28 kPa). Pregerminated plants of *P. virgatum* and *R. hirta* were transplanted into the pots (one plant per pot) and grown for 48 days in a controlled growth chamber. Additionally, three unplanted control pots were prepared and maintained under the same conditions as the planted pots. Growth chamber conditions were kept constant with 14 hr photoperiod, 400 µmol m^−2^ s^−1^ photosynthetically active radiation, and 25°C and 20°C day and night temperature. An additional 200 ml of the nutrient solution was added to the pots on the 10^th^ and 20^th^ days after transplanting. The pots were watered every other day with the amount of water sufficient to maintain 30% water content added from the bottom, which was covered by a 30-μm nylon mesh.

**Figure 1:**
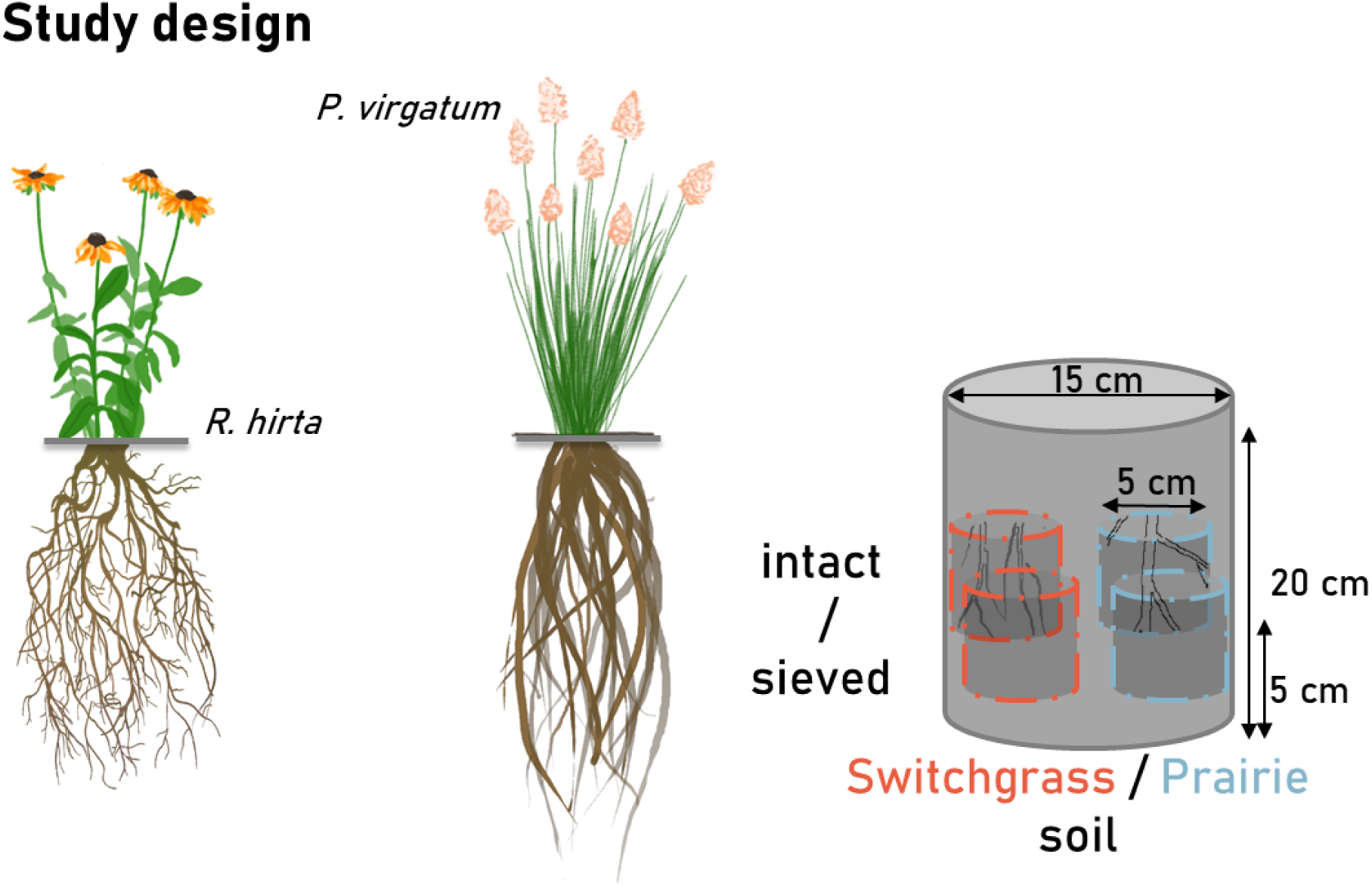
Outline of the study’s experimental design. *R. hirta* and *P. virgatum* were planted into containers filled with 2-mm sieved soil and holding perforated ingrowth cores with soil of four contrasting pore structure characteristics, namely: intact and sieved soils of prairie and switchgrass origin. To ensure that the intact and sieved cores differed from each other only in terms of pore structure, the sieved soil was packed to the same bulk density as the intact cores and all >2 mm inclusions, such as stones and large root residues were incorporated back into the sieved soil prior to packing.

Each core was subjected to X-ray µCT before and after the plant growth experiment. Details on X-ray µCT scanning and image processing are reported in Lucas et al. (2023b) and presented in the Appendix (Method S1). Double scanning enabled us to describe root growth patterns as a function of soil origin and structure as well as to quantify the proportions of roots growing into soil regions with disparate pore structures. Table 1 summarizes plant growth parameters as well as the share of roots grown into soil regions with different pore structures, namely, into soil dominated by < 0.04 mm diameter pores (soil matrix), into macropores (> 0.04 mm diameter pores), and into biopores (as determined by Lucas et al 2023). The latter include all macropores of biological origin and thus of cylindrical shape.

**Table 1:**
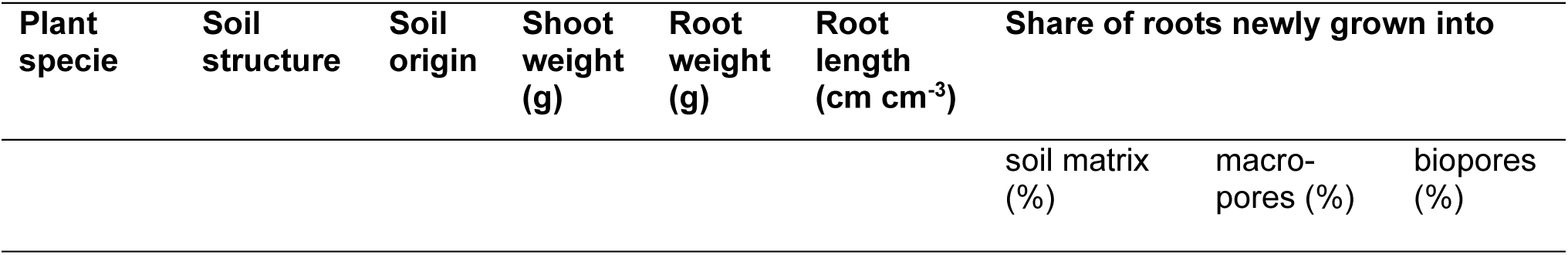
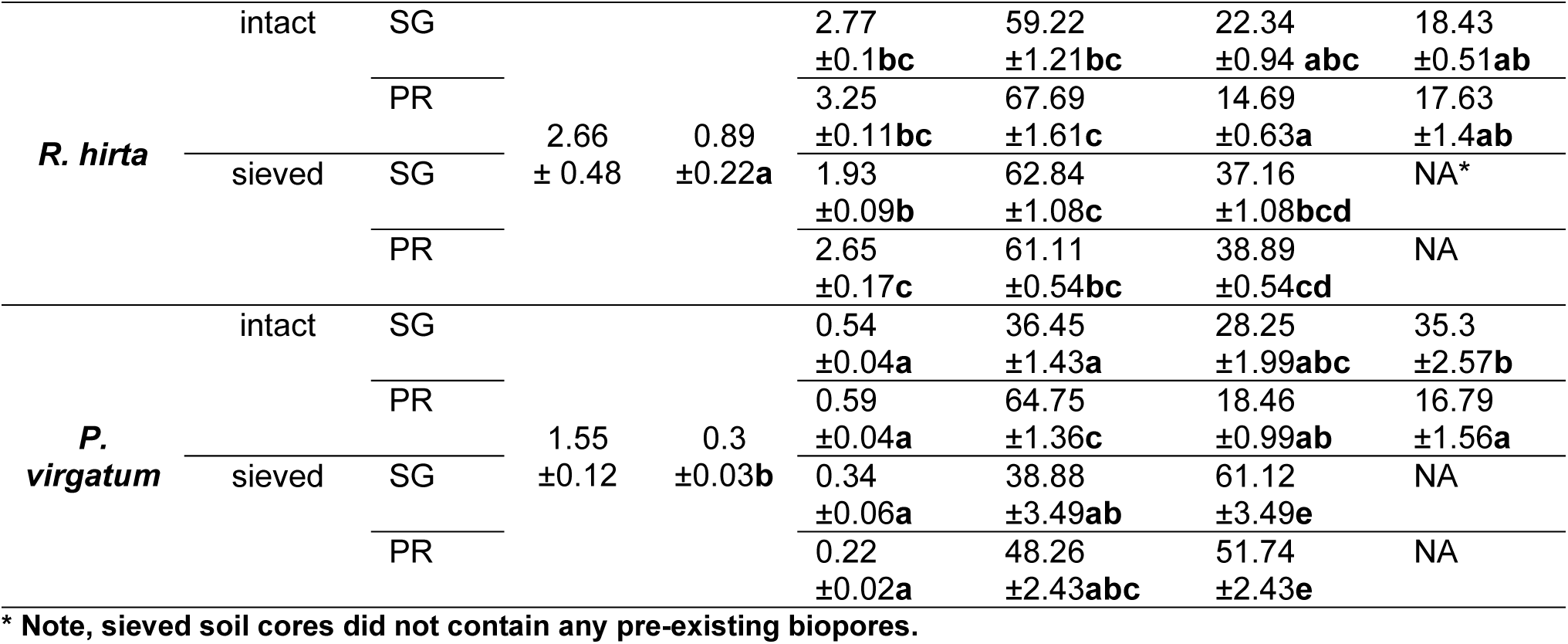
Summary of plant and root growth pattern characteristics (reported in Lucas *et al*., 2023b). Depicted are means and standard errors of shoot and root weights (n=6) for *R. hirta* and *P. virgatum* and their root length in the soils of the two studied origins (Switchgrass = SG and Prairie = PR soil) and structures (intact and sieved). In addition, the share of roots grown into soil matrix (< 0.04 mm diameter pores), macropores (> 0.04 mm diameter), and biopores is reported as % of the total root volume. Different letters mark significant differences between the studied plants for shoot and root weights as well as among the studied plant, origin, and structure combinations for the rest of the data (p-value<0.05).

The plants were labelled with ^14^C-CO_2_ according to Santiago et al. (2021) using LI-COR 6800 portable photosynthesis system (Li-Cor, Lincoln, NE, USA) with an 3×3 cm chamber to feed plant leaf’s with CO_2_. A constant flow from a pressurized ^14^CO_2_ tank ensured a concentration of 420 µmol mol⁻¹ of labelled CO_2_ in the chamber. To ensure a distribution of ^14^C in the whole root system (Pausch & Kuzyakov, 2011) as well as sufficient amounts of detectable ^14^C in the rhizosphere, the plants were labelled at 11th and 3rd day before harvest under the same environmental conditions as set in the growth chamber.

### 2.2 Analyses of rhizosheath and rhizodeposition

During the harvest, the pots were disassembled, and the cores were removed. The cores were cut open, and roots were meticulously extracted from the soil using tweezers. The roots were gently shaken to detach loosely adhering soil, and the root-adhering soil, termed rhizosheath, was carefully removed with small brushes. The rhizosheath soil was then oven-dried at 60°C to determine its dry weight. Subsequently, the dried rhizosheath soil was combusted in an oxidizer (PerkinElmer®, Boston, MA, Model A307), and the released ^14^CO_2_ was trapped in 20 ml of scintillation cocktail (Carbo-Sorb E:Permafluor® E+, 1:1 [v/v], PerkinElmer, Groningen, Netherlands). The radioactivity of the rhizosheath was measured using a liquid scintillation counter (PerkinElmer® Tri-Carb 4910TR, Waltham, MA).

### 2.3 Image analysis

Pore size distributions (PSD) were calculated from the segmented pore/solid binary images. PSD was determined in Fiji (Schindelin *et al*., 2012; Rueden *et al*., 2017) using the local thickness method (Hildebrand & Rüegsegger, 1997). Images from after the experiment contained both newly grown roots and old roots from the field. To differentiate new from old roots, all roots were segmented in images before and after the experiment and subtracted from each other to create a mask of roots grown during the experiment. Roots and biopores were segmented as described in Methods S1. Root growth preferences were analyzed by using the newly grown root mask on the segmented image from before the experiment (labeled by macropores, matrix, biopores, and roots, Fig. S1) to calculate the volume percentage of each structure encountered by the root.

Rhizosphere macroporosity gradients were calculated using the Euclidean Distance Transform (EDT) in Fiji, following Lucas et al. (2019) and as depicted in Fig. 2A. Besides the overall macroporosity (pores > 0.04 mm), we focused on the formation of narrow pores (0.04-0.15 mm). While the lower boundary of 0.04 mm is set due to the image resolution, pores with a diameter of more than a few tens of micrometres up to 0.15 mm offer ideal conditions for microbial activity (Ruamps *et al*., 2013; Kravchenko *et al*., 2019; Li *et al*., 2024). The EDT computes the Euclidean distance to the nearest root for every voxel in the 3D images. The EDT of the newly developed roots was combined with the binary pore images and the PSD images from before and after plant growth to calculate gradients of all macropores and narrow pores around the roots, or more precisely their local percentage to the total local soil volume. Note that in this analysis biopores are included, being only a subset (in shape) of macropores of different sizes. The resulting pore gradients from before and after root growth were used to differentiate between changes directly induced by the roots, i.e., root-driven changes, and macroporosity gradients resulting from inherent root growth preferences (Fig. 2B). The changes in macroporosity gradients due to root activity were calculated by subtracting the local macroporosity, i.e. the macroporosity at a given distance to the root, before root growth from that after root growth; these will be referred to as ‘root-induced changes’. The macroporosity gradients resulting from root growth preferences were calculated by subtracting the mean total macroporosity of the sample from the local macroporosity and will be referred to as the ‘root growth preference effect’.

**Figure 2:**
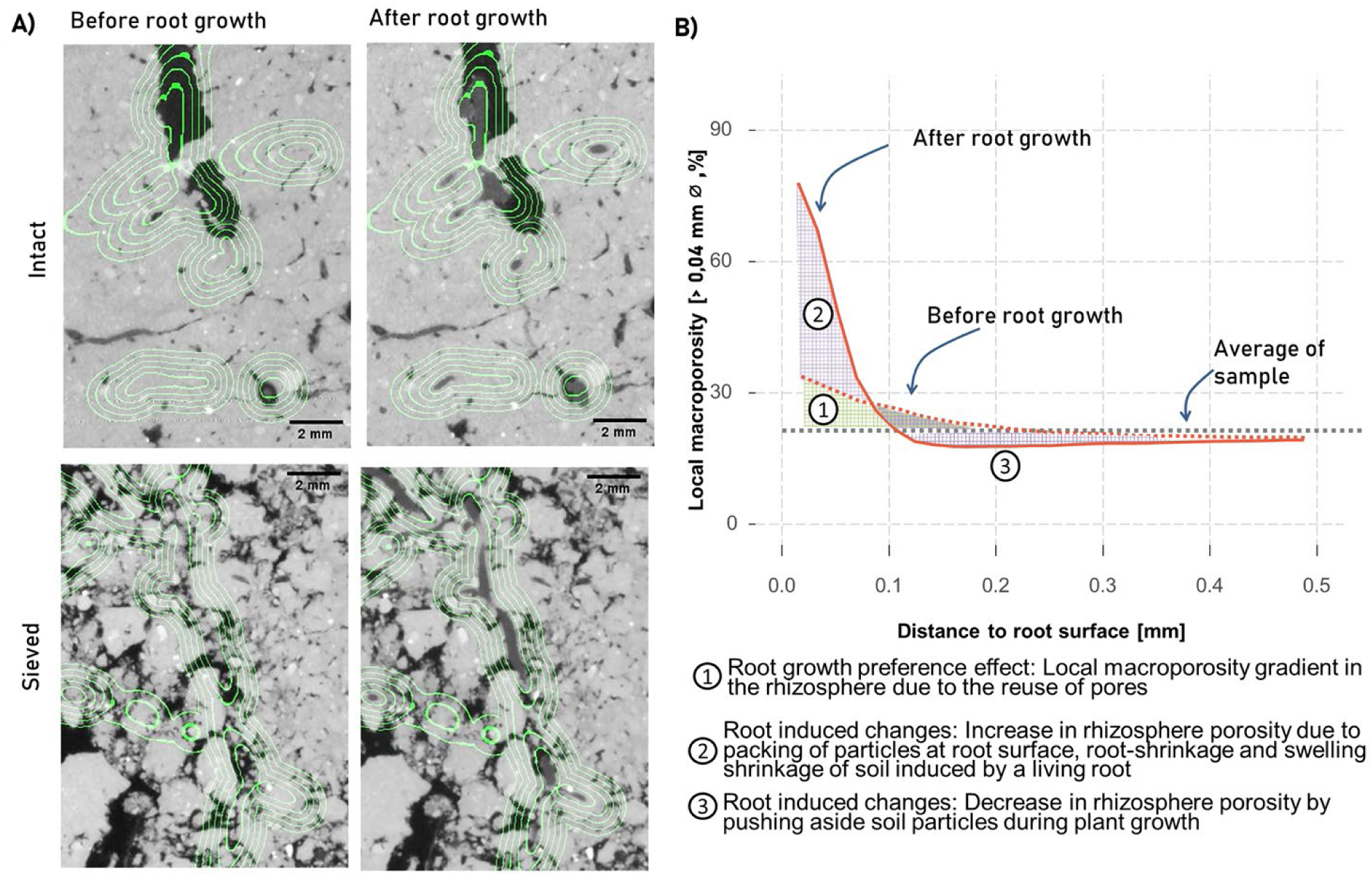
Methodological approach for image analysis developed for quantification of root-induced changes and the root growth preference effect. A) Small image sections of X-ray CT scans illustrating the approach. The ingrowth cores were scanned twice to derive macroporosity profiles before and after root growth in all sieved and intact samples. Green lines mark examples of specific distances (0.2,0.4, 0.6, 0.8, 1.0 mm) along these profiles to the roots grown within the experiment. Note that these distances are calculated based on 3D images (Fig. S1), thus may appear as somewhat off when overlaying a single 2D slice in this illustration. B) Schematic representation of the approach used to evaluate macroporosity gradients around roots using X-ray µCT. The soil induced root growth preferences effect on the macroporosity gradients (1) is the difference between the average macroporosity of the core and the local macroporosity around the root. The changes induced by plants (2,3) are the difference between the macroporosity profiles before and after root growth.

These values were calculated for a range of distance increments from the root surface aimed at characterizing the spatial distribution patterns within the most active part of the rhizosphere. Overall, the size of the rhizosphere, that is the soil directly influenced by the root, largely depends on the process being investigated and a variety of soil and root characteristics as well as measurement tools, and can range from 0.3 to 5 mm distances from root surfaces (Hinsinger *et al*., 2009; Marschner, 2012; Kuzyakov & Razavi, 2019). Here we focused on the 0-1 mm interval, a conservative estimate representing the zone most strongly affected by root influences on pore structure. We explored that interval in 0.2 mm increments, where each increment encompasses ∼ ten image voxels, a number sufficient to provide a reliable estimate of the image-derived characteristics (Vogel *et al*., 2010). While the voxel size of 0.0182 mm allowed for sufficient segmentation of roots - whose diameters span several voxels (Lucas *et al*., 2022, 2023b,a) – partial volume effects prevent a perfectly sharp distinction of the root boundary. To ensure the accuracy of our statistical analysis, we excluded the border voxel, as it may not precisely represent the actual root boundary. However, larger errors spanning multiple voxels can be ruled out, as they would be evident during visual inspections of the segmentation results.

As grey values from X-ray µCT images depend on electron density, they were previously used as a reliable indicator for local gradients in soil density (Lucas *et al*., 2019; Phalempin *et al*., 2023). Thus, we calculated gradients in grey values around roots after the plant growth in the same way as done for macropores, i.e. using EDTs. The gradients in grey values were used to calculate the mean grey values in the rhizosphere, named hereafter µCT-derived rhizosphere density, to associate rhizosphere density with conventionally measured rhizosheath mass (see Statistics section below).

### 2.4 Statistics

We tested the effects of plant species (*P. virgatum* vs. *R. hirta*), soil structure (sieved vs. intact), soil origin (switchgrass soil vs. prairie soil) and the type of process leading to changes in local macroporosity in the rhizosphere (root-induced changes vs. root growth preferences) on 1) macroporosity changes and 2) narrow pore changes as a function of the distance from the roots. The distance was treated as a categorical variable with 5 levels representing 0.2 mm distance increments within the target 0-1 mm distance from the roots (See Table S1 for detailed model structure). The selected distance from the root to be included in this analysis, i.e., 0-1 mm, is consistent with the portion of the rhizosphere that is commonly found as the most affected by the roots’ influences (Lucas *et al*., 2019; Phalempin *et al*., 2021), while the 0.2 mm bin size enabled capturing the spatial gradient with sufficient number of data points within each bin. Note that we verified that the conclusions remained unchanged when the distance was binned at 0.1 mm instead of 0.2 mm (data not shown). The statistical model (model 1) included these five factors and their interactions as fixed effects. Random effects included the planted pots, used as an error term for testing the plant species effect, and the ingrowth cores nested within the pots, used as an error term for testing the effects of soil structure and origin.

The statistical model (model 2) for rhizosheath dry weight data consisted of the fixed effects of plant species (*P. virgatum* vs. *R. hirta*), soil structure (sieved vs. intact), soil origin (switchgrass soil vs. prairie soil) and random effects of the planted pots, used as an error term for testing the plant species effect, and the ingrowth cores nested within the pots, used as an error term for testing the effects of soil structure and origin. The models were fitted using *nlme* function of the *lme4* package (Bates *et al*., 2015) of R (R Core Team, 2023, V. 4.1.1).

Analysis of the residuals was performed for all studied response variables of model 1 to assure that there were no model overfitting issues (Fig. S2). The assumptions of normality and homogeneity of variances were assessed using normal probability plots of the residuals and Levene’s tests for equal variances. To achieve a normal distribution of the model residues for the model 1, the function “Gaussianize” was used (Goerg, 2015), while the rhizosheath weight for model 2 was log-transformed. When the equal variance assumption was violated, the unequal variance model were fitted using the *nlme* package in R (Pinheiro & Bates, 2000; Pinheiro *et al*., 2024). Unequal variances for distance increments were invariably required in model 1 since both the mean values of the studied responses and their variability were markedly higher in close proximity to the roots, while decreasing with the distance.

Since interactions among the studied factors were statistically significant (p-value < 0.05), for model 1 slicing of the interactions, aka simple effect F-tests, were conducted using the *emmeans* package (Lenth, 2024) to detect differences between the two processes at various distances. For model 2, pairwise comparisons of the estimated marginal means were conducted using the cld-function of the *emmeans* package.

To explore the associations between local root-induced changes in the rhizosphere and the overall microporosity we used linear regression analyses. To confirm that the root-growth preference effect is indeed largely driven by root growth into macropores vs. root growth into the soil matrix, the linear regression analyses were performed between numbers of roots grown into the soil matrix (as calculated in Lucas *et al*., 2023b) and macroporosity changes in the rhizosphere. This was done not only for the whole rhizosphere (i.e., 1 mm distance to the root), but also for the first 0.2 mm of the rhizosphere. The latter was regarded as necessary based on distinct patterns of the root induced-changes within the first 0.2 mm of the rhizosphere detected using the mentioned above model 1 and 2 results. To analyze properties associated with rhizosheath dry weights, linear regressions of the shares of roots grown into soil regions of different structures (i.e., soil matrix, macropores, biopores) on the rhizosheath dry weight were computed. In addition, we conducted linear regression analysis of the associations between the rhizosheath dry weight and the µCT-derived rhizosphere density (image grey values. Spearman correlation coefficients were calculated to explore the association between rhizosheath weight and µCT-derived rhizosphere densities.

## 3 Results

### 3.1 Rhizosphere macroporosity gradients

Gradients in macroporosity at the locations later populated by roots were already observed prior to root growth, and across all investigated soil origins and structures the macroporosity near these locations was higher compared to the average macroporosity (Fig. 3). Root growth further modified these gradients, particularly within the <0.2 mm distance from the root surface, where a significant increase in macroporosity was observed for both plants across all four soil origins and structures.

**Figure 3:**
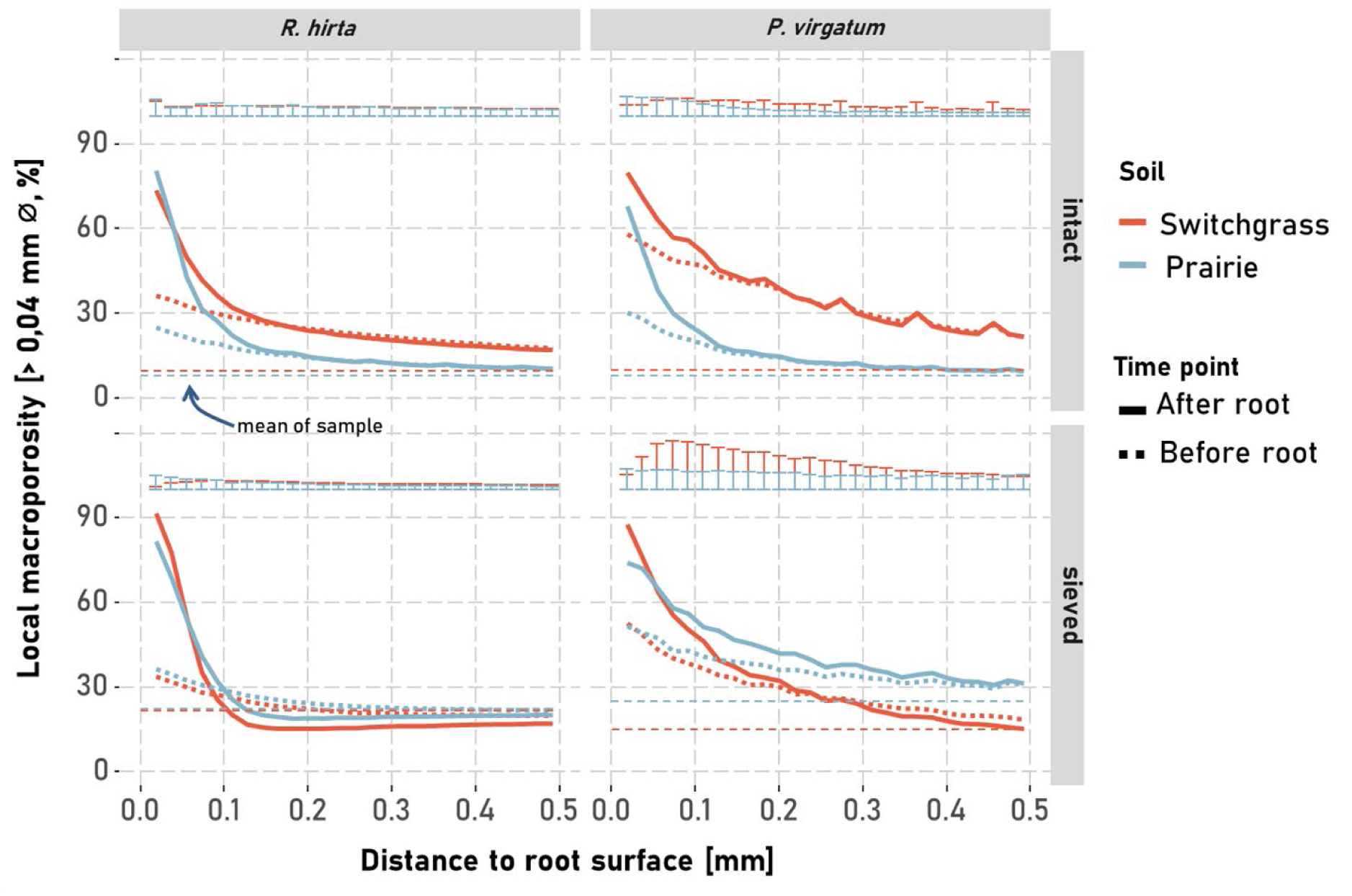
Gradients of the local macroporosity (pores >0.04 mm) with distance to the root surface of *Panicum virgatum* and *Rudbeckia hirta* when grown in sieved or intact soil of prairie or switchgrass origin. Gradients of macropores computed before (dotted line) and after root development are shown, along with average total macroporosity (dashed line). Error bars represent the standard errors of the local macroporosity means after root growth (n=6).

These effects of root growth preference and root-induced changes on macroporosity were strongly influenced by plant type and soil structure but not by soil type (Table S1). Root growth preferences consistently increased rhizosphere macroporosity near the root surface, gradually tapering to background levels with distance (Fig. 4). At most distances, this effect exceeded that of root-induced changes. However, within 0.2 mm of the root, both processes increased macroporosity substantially (5–30%). Root-induced changes within this range were similar for *R. hirta* and *P. virgatum*, regardless of soil structure or origin, and increased macroporosity by 5–10%. Beyond 0.2 mm, this increase in macroporosity declined rapidly. In sieved soils macroporosity even decreased through root-induced changes by *R. hirta* at distances >0.2 mm. In contrast, root growth preferences showed greater variability within 0.2 mm. For example, *R. hirta* increased macroporosity by ∼5% in sieved soil, while *P. virgatum* caused a >30% increase in switchgrass soil—more than twice the effect seen in prairie soil.

**Figure 4:**
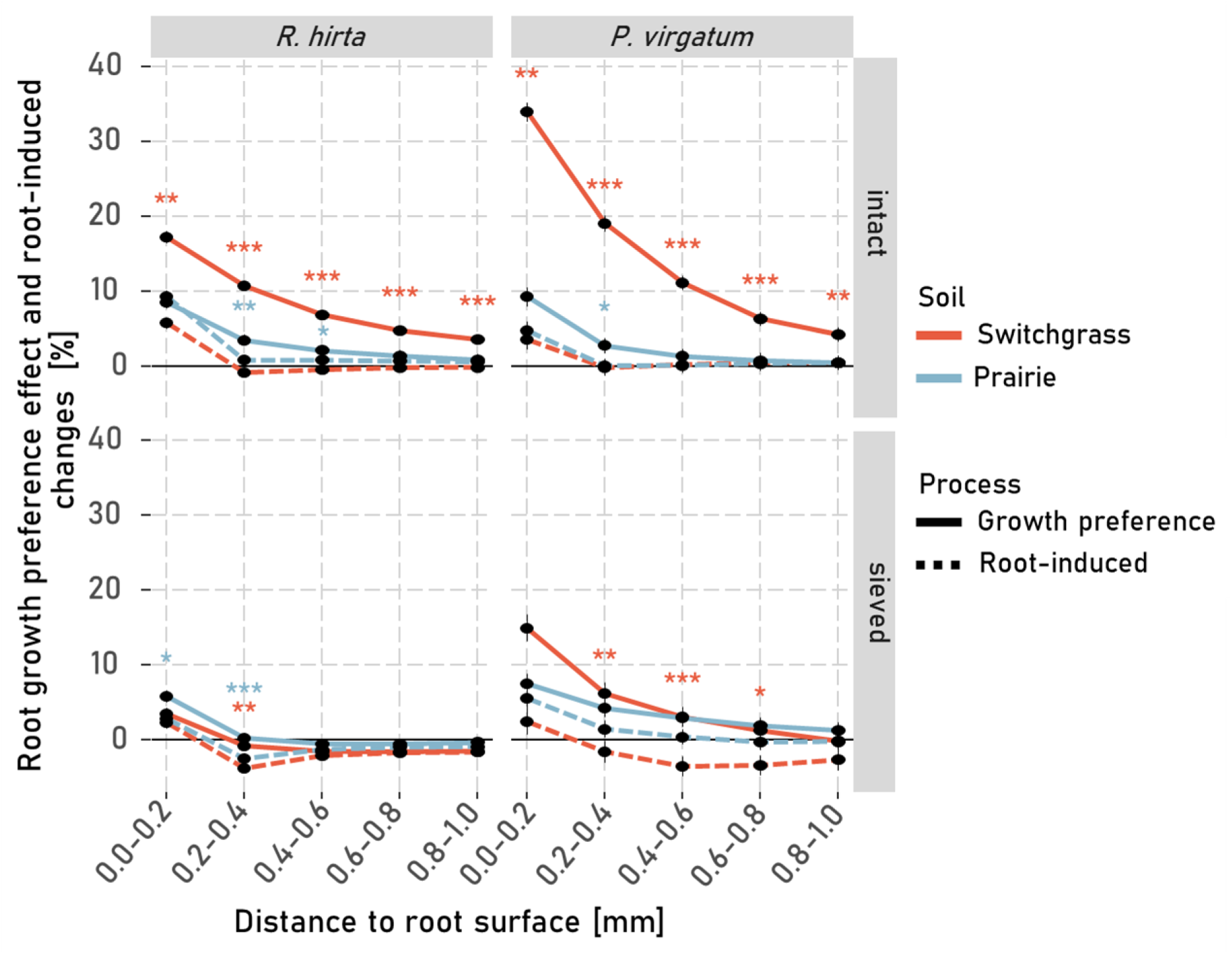
Local macroporosity changes in the rhizosphere due to the root growth preference effect (solid line) and root-induced changes (dashed line) of *P. virgatum* and *R. hirta* when grown in sieved or intact soil of prairie or switchgrass origin. Local macroporosity changes are calculated as described in Fig. 2. Asterisks indicate significant differences between the root-induced changes and the changes resulting from root growth preferences at a given distance. For p-values < 0.05, < 0.01 and < 0.001 *, ** and *** are displayed. Whiskers represent the standard errors of the mean (n=6).

As expected, the local macroporosity changes from the root growth preference effect was strongly negatively correlated with the proportion of roots grown into the soil matrix, i.e., the soil areas lacking macropores, at both <0.2 mm (R²=0.54, p-value < 0.001, Fig. S3A) and <1 mm (R²=0.26, p-value < 0.001, Fig. S3B) distances. In contrast, root-induced changes on local macroporosity at <0.2 mm were positively associated with the proportion of roots grown into the soil matrix (R²=0.14, p-value = 0.012, Fig. S3C), but not for a larger rhizosphere volume, i.e. <1 mm distance to the root (R² = 0.03, p-value = 0.30).

### 3.2 Effects on narrow pores

The volume of the narrow pores (0.04-0.15 mm diameter) was generally low, especially in the cores with intact soil structure (Fig. S4). Consequently, the root growth preference effect on such pores was rather minor (< 3%, Fig. 5) for both *P. virgatum* and *R. hirta*. Interestingly, root-induced changes led to similar patterns for the narrow pores (Fig. S4) as compared to all macropores (Fig. 3), with a stark increase in these pore sizes close to the root surface. There were hardly any changes in intact soil compared to compaction in sieved soils at >0.2 mm distances (Fig. 5). However, note that at distances beyond 0.4 mm from the root, porosity differences remained statistically detectable for *R. hirta* growing into intact soil but were consistently small (<1%; Fig. 5).

**Figure 5:**
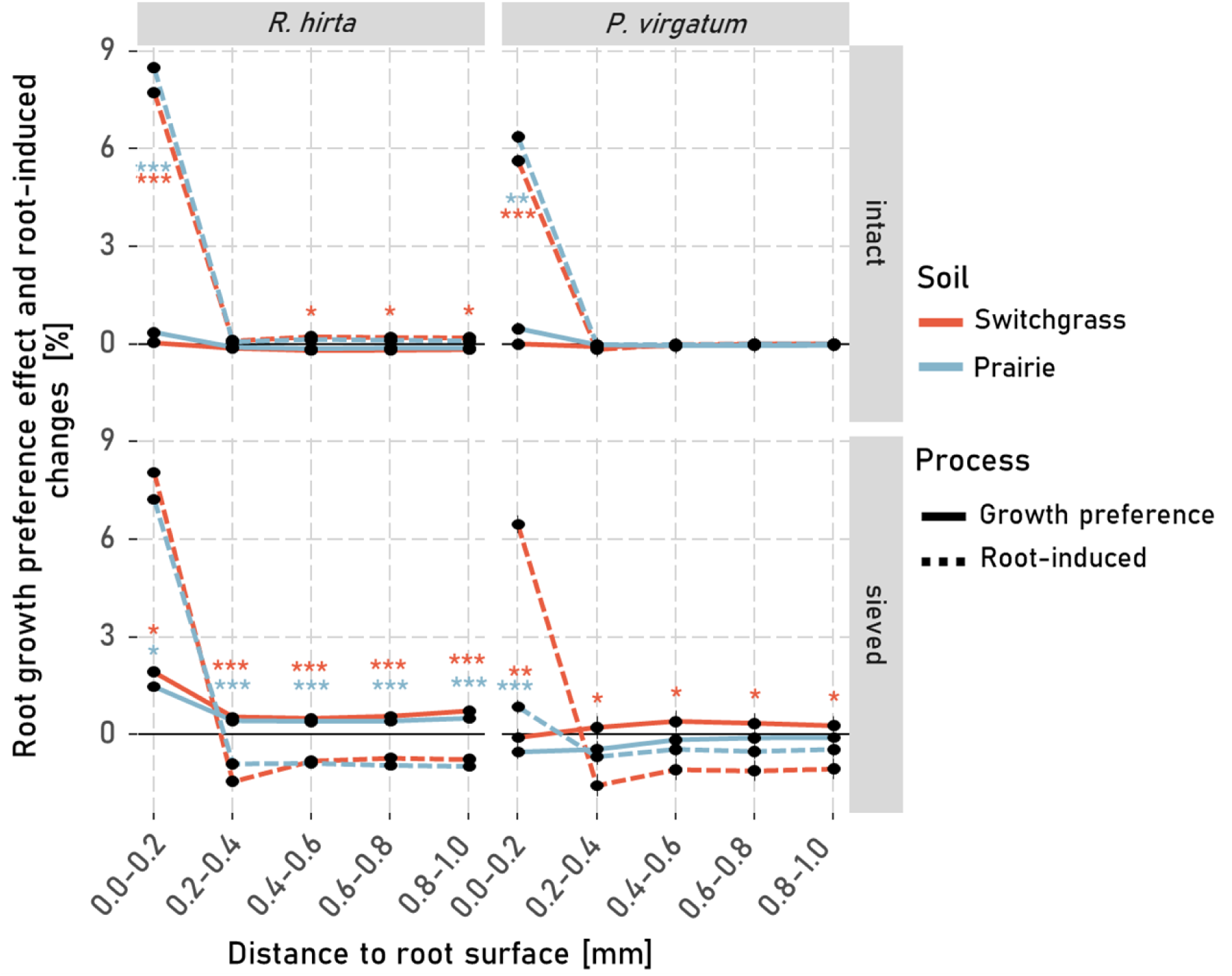
Local changes in narrow pores within the rhizosphere due to the root growth preference effect (solid lines) and root-induced changes (dashed lines) of *P. virgatum* and *R. hirta* when grown in sieved or intact soil of prairie or switchgrass origin. Local porosity changes are calculated as described in Fig. 2. Asterisks indicate significant differences between the root-induced changes and the changes resulting from root growth preferences at a given distance. For p-values < 0.05, < 0.01 and < 0.001 *, ** and *** are displayed. Whiskers represent the standard errors of the mean (n=6).

### 3.3 Impact of Root-Induced Changes on macroporosity

There were no significant differences in total macroporosity between the two plant species and the unplanted controls in any of the investigated settings (Fig. 6A). However, while macroporosity in intact soil cores remained unchanged (Switchgrass soil) or numerically increased (Prairie soil), all sieved soil cores exhibited a reduction in macroporosity during plant root growth. The linear-regression between root-induced changes in the rhizosphere and changes in the total macroporosity of the samples was significant and close to the 1:1-line (Fig. 6B, R^2^=0.81, p-value <0.001), whereas the root growth preference effect was not associated with the changes in total macroporosity (Fig. S5).

**Figure 6:**
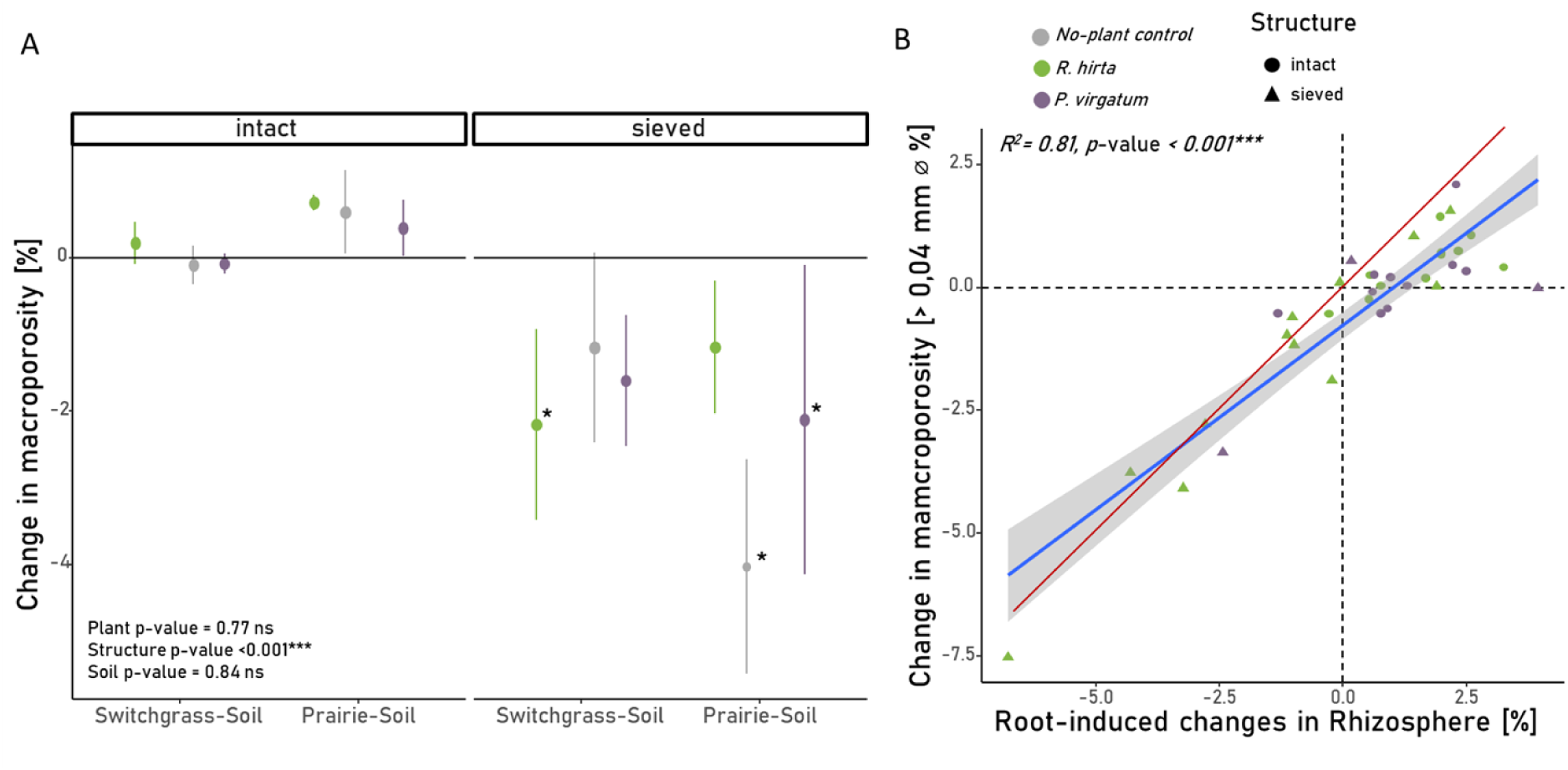
A) Changes in total macroporosity in cores from *P. virgatum*, *R. hirta*, and the unplanted control for the sieved and intact soils of prairie or switchgrass origin. There were no significant differences between plant species and soil origin combinations within each structure, but the effect of structure was significant, as indicated in the lower left part of the plot (p-value < 0.001). Stars indicate significant differences to zero (p-value < 0.05). b) Association of root-induced changes of local macroporosity in the rhizosphere (< 1 mm distance to the root) with the change in total macroporosity after plant growth. Red-line represents 1:1-line. Samples with root length <10 cm per core were excluded as their influence on total macroporosity was considered negligible. Statistical significance is marked by ns, *, **, and *** for p-values > 0.1, < 0.05, < 0.01, and < 0.001, respectively

### 3.4 Rhizosheath formation and it’s relation to rhizosphere properties

Rhizosheath dry weight exhibited high variability, with significantly lower values for *R. hirta* grown in intact Prairie soil and sieved Switchgrass soil compared to *P. virgatum* grown in sieved Switchgrass soil (Fig. 7A). The rhizosphere density gradients showed similar patterns as observed for the gradients of the macroporosity only, with the highest decline in density close to the root surface (Fig. S6). Surprisingly, there was no correlation between the µCT-derived rhizosphere density and rhizosheath dry weight for *P. virgatum* for any distance increment (Fig. 7B). In contrast, there was an association of µCT-derived rhizosphere density and rhizosheath mass for *R. hirta* especially if the volume >1mm distance is included the estimation of the rhizosphere density. This association, however, was negative. However, for both plants the rhizosheath weight was positively correlated with plant-derived C, i.e. ^14^C activity, in it (Fig. 7C), and with the proportion of roots growing into macropores (Fig. 7D). There were no significant associations between the proportion of roots in biopores or in the soil matrix and the rhizosheath dry weight (Fig. S7a,b).

**Figure 7:**
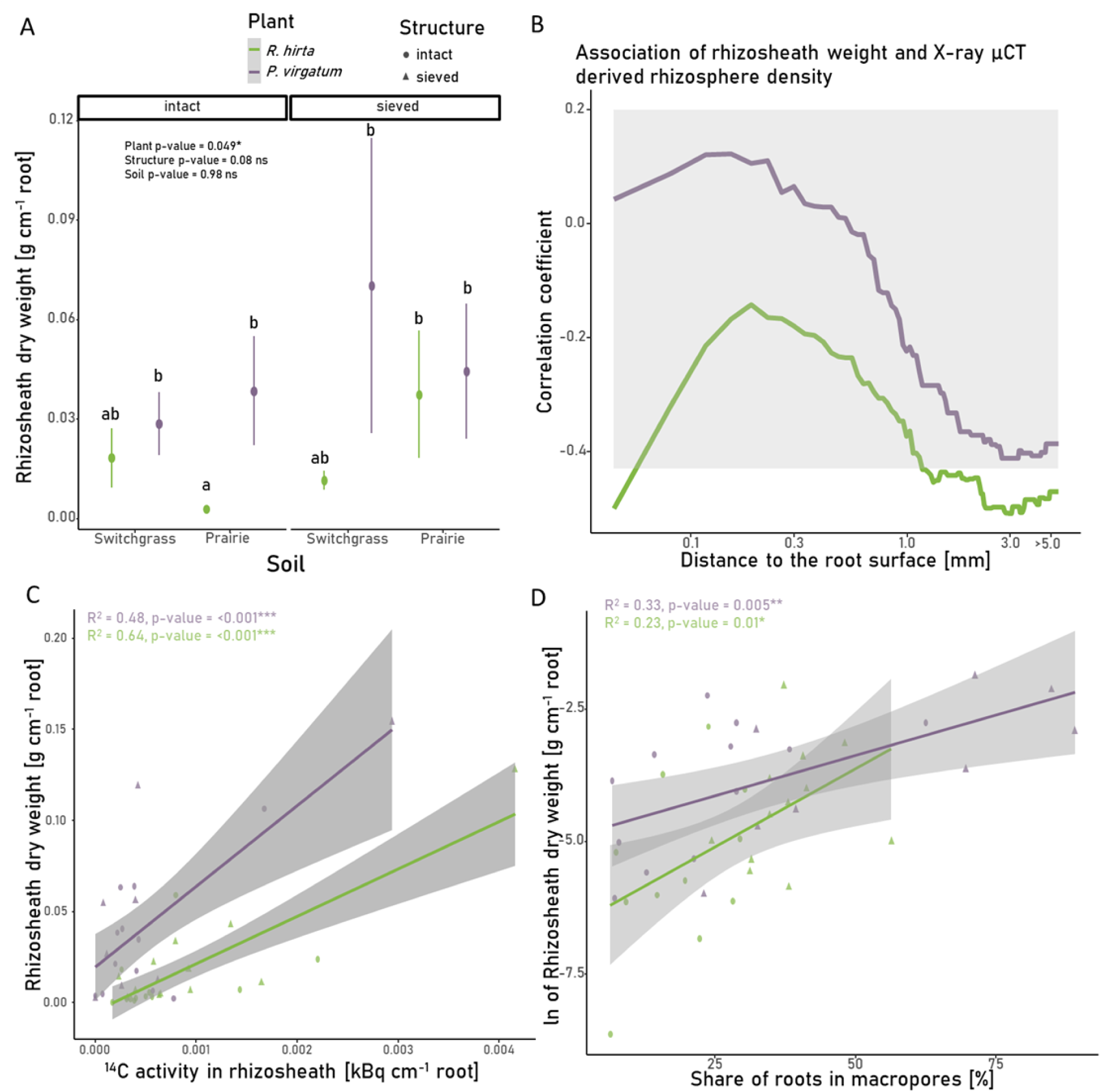
Rhizosheath development of *P. virgatum* and *R. hirta*. a) Rhizosheath weight of *P. virgatum* and *R.hirta* roots grown into intact and sieved soil of the two studied origins. Shown are mean values and standard errors. The letters indicate significant differences among the plant, structure, and soil origin combinations (p-value < 0.05). b) Spearman-correlation coefficients for rhizosheath dry weights and X-ray CT derived rhizosphere density, where the latter are estimated as the mean density of the soil starting at the root and ending at the depicted distance. Grey area marks non-significant correlations (p-values >0.05). Note the non-linear x-axis. c) Plant-derived C, represented by ^14^C activity, in the rhizosheath plotted vs. the rhizosheath weight. d) Share of roots grown into macropores plotted vs. the rhizosheath weights. In b)-d) dots are the observed data and lines are the linear regression models fitted to them. Statistical significance is marked by ns, *, **, and *** for p-values > 0.1, < 0.05, < 0.01, and < 0.001, respectively

## 4 Discussion

### 4.1 Root-induced vs. root growth preference-driven porosity gradients in the rhizosphere

Our findings from the sieved soil align well with previous studies exploring rhizosphere characteristics in similarly finely sieved and repacked soil settings (Helliwell *et al*., 2017; Koebernick *et al*., 2017, 2019; Phalempin *et al*., 2021). However, by using repeated X-ray µCT imaging, we were able to describe rhizosphere macroporosity gradients forming upon new root growth not only in sieved but also in natural, intact field soil. The approach made this study the first to distinguish between root-induced changes in rhizosphere macroporosity and the macroporosity gradients resulting from the root growth preference effect.

The results partially supported our first hypothesis indicating that root-induced changes contributed less to the formation of rhizosphere porosity gradients than the influence of root growth preferences. However, the relative contributions of the two mechanisms depended on the position within the rhizosphere (Fig. 4) and on the magnitude of root growth into the soil matrix (Fig. S3). Close to the root-interface, i.e., at <0.2 mm distance, direct influences of roots on the rhizosphere macroporosity were comparable (in sieved soil) or smaller (in intact switchgrass soil) than the root growth preference effect (Fig. 4). At distances >0.4 mm, root-induced changes were generally minor whereas the root-growth preference effect especially in intact soils still substantially influenced local macroporosity in the rhizosphere.

Root-induced changes are a combined result of enhanced shrinkage and swelling of the soil, rhizodeposition, root hair growth and root shrinkage (Carminati *et al*., 2013; Koebernick *et al*., 2017; Helliwell *et al*., 2019). When a larger proportion of roots grew into the soil matrix, root-induced changes within <0.2 mm from the root surface were the highest (Fig. S3C). These changes appeared to be primarily driven by the formation of pores of a specific size, namely narrow pores (Fig. 5). This suggests that geometric factors strongly influence the root-induced changes at the root-soil interface for roots growing into the soil matrix (Koebernick *et al*., 2019). When roots push aside and rearrange soil upon their growth through the soil matrix, the rearrangement and packing structure of the relatively spherical soil particles along the flat root surface naturally lead to increased porosity at the immediate root interface—a phenomenon described as the surface-wall effect (Koebernick *et al*., 2019; Lucas *et al*., 2019). Moreover, when roots grow into the soil matrix and rearrange the pore system, they may lead to rhizosphere compaction. Consistent with this notion and in line with previous studies (Koebernick *et al*., 2019; Lucas *et al*., 2019; Phalempin *et al*., 2021), we observed that macropores within 0.2-0.4 distance were compressed by root growth. In general, soil compression mainly reduces large structural macropores, with a lesser impact on meso- and micropores within the soil matrix (Kutilek *et al*., 2006). However, macropores that are as large as, or larger than, the roots themselves can be reused by roots and are therefore typically not compacted during root growth (Lucas *et al*., 2022). Thus, it is primarily the volume of narrow pores that is affected by the compaction. Indeed, in the sieved soil, these narrow pores were the ones primarily compacted by plant roots (Fig. 5 bottom panels). This was especially true for the roots of *R. hirta*, even though their root diameter was smaller when grown into sieved soil compared to the roots in intact samples (Fig. 3C in Lucas et al 2023). In contrast, in intact field soils where many roots grew into existing biopores (Lucas et al. 2023) and where the narrow pores were less abundant, root-induced changes on macropores in the rhizosphere by roots were minimal (Fig. 4 and 5 top panels). This explains why a compacted rhizosphere zone is typically observed in finely sieved soils with a large amount of narrow pores (Koebernick *et al*., 2019; Lucas *et al*., 2019) but is absent in loosely packed or coarsely structured soils with many large macropores, as well as in field soils with biopores (Lucas *et al*., 2019; Phalempin *et al*., 2021). The lack of a compaction zone in intact samples is, therefore, a result of fewer narrow pores and of more well-connected biopores available for reuse. The strong negative correlation between the root growth preference effect and the proportion of roots growing in the soil matrix (Fig. S3B) emphasizes that the roots’ preference to grow into more porous and macropore abundant areas was the primary driver of the observed local macroporosity gradients in the rhizosphere. Thus, while previous studies suggested that coarse root systems induce higher local compaction (Dexter, 1987; Lucas *et al*., 2019), our results demonstrate that even for coarse rooted plants, such as *P.virgatum*, the reuse of existing pores can be the main factor shaping rhizosphere macroporosity (Fig. 4, top right panel). These findings underscore the importance of the interactions between roots and the inherent pore structure of the intact soil in defining rhizosphere pore characteristics (Lucas *et al*., 2019; Phalempin *et al*., 2021).

The macroporosity gradients in the rhizosphere show a strikingly consistent pattern across different soil structures, particularly for narrow pores (Fig. S4). In both intact and sieved soils, the volume of such pores increases sharply close to the root surface tapering down to very low values at 0.2–0.4 mm. In intact soils, such low values result from the low number of narrow pores in the bulk soil, while in sieved soils, it is primarily due to root-induced compaction (Fig. 5). Although intact and sieved soils initially displayed different porosity patterns, they have now converged to a remarkably similar profile, likely due to the interplay between root growth preferences and root-induced changes that counterbalance each other’s effects (Fig. S3A, C).This finding aligns with the concept of a self-organizing rhizosphere, which optimizes conditions for microbial habitats and water movement (Rabbi *et al*., 2018b). Young & Crawford (2004) introduced the idea of a self-organizing soil-microbe complex as the interconnected relationship between soil’s physical structure (e.g., its pore network) and microbial life. As microbes break down organic matter and interact with root carbon inputs, they can reorganize the soil’s pore structure. This reorganization affects how water, nutrients, and air move through the soil, in turn affecting the plants and the microbes themselves. The two systems - biological and physical - are therefore intertwined, constantly influencing each other in the rhizosphere environment (Young & Crawford, 2004; Rabbi *et al*., 2018b). Our results demonstrate that a joint impact of the root growth preference effect and root-induced changes modifies rhizosphere pore structure by increasing proportions of narrow (0.04-0.15 mm diameter) pores in close proximity to the roots, that is, exactly in the locations where root exudates and rhizodeposits are released. Thus, soil-root-microbe interface provides an optimal environment for microbial activity, making root surfaces and their immediate vicinity particularly active microbial hotspots (Schmidt *et al*., 2018; Mueller *et al*., 2024). In addition, the reduction in narrow pores in the subsequent rhizosphere (0.2 to 0.4 mm) can lead to longer periods of lower transpiration during dry conditions, thus potentially protecting the plant from rapid dehydration (Landl *et al*., 2021). In summary, the interactions between the root growth preference effect and root-induced changes play a key role in the self-organization of the rhizosphere, establishing favorable physical conditions for nutrient acquisition and water uptake by plants across both sieved and intact soil structures (Rabbi *et al*., 2018b; van Veelen *et al*., 2019; Landl *et al*., 2021).

### 4.2 Changes in total macroporosity

Consistent with our expectations, minimal changes in the total macropororsity were observed in intact soils, with no differences from unplanted controls (Fig. 6A). Past studies reported high variability in plant induced changes in macroporosity, depending on the plant type and soil pore structure (Bodner *et al*., 2014; Lucas *et al*., 2022). Yet, consistent with our results, when there is a well-developed macropore system, plants seem to reuse and rearrange the existing pore space without affecting the pore structure (Lucas, 2020; Lucas *et al*., 2022).

The total macroporosity in the sieved samples decreased upon root growth (Fig. 6A), and the reduction in the macroporosity in the rhizosphere from the roots apparently reduced the total macroporosity (Fig. 6B). Note, however, that macroporosity in sieved Prairie soil also decreased in the no-plant controls (Fig. 6A). The decrease in porosity in no-plant controls suggests a settling of the sieved pore structure. The reduction in local macroporosity observed at larger distances from roots (>1 mm) in sieved soil may therefore partially explained through an overall reduction in macroporosity without the action of plants (Fig. 4, bottom panels). In addition, even in intact soil at distances greater than 0.4 mm, significant root-induced changes were observed (Fig. 5, top left panel); however, their magnitude was very small, typically <1%. Such small changes could also result from the degradation of particulate organic matter over the experimental period, leading to the formation of new small pores rather than direct root activity. Therefore, these effects at higher distances in both intact and sieved soils are unlikely to be of physiological or physical relevance.

### 4.3 Rhizosheath Development

Our second hypothesis, proposing that rhizosheath forms as roots penetrate the soil matrix and thereby increase soil–root contact, densify the rhizosphere structure, and enhance root-derived carbon inputs, was only partially supported by the data. We observed strong positive associations between the rhizosheath mass and the quantities of root-derived carbon in it (Fig. 7C). However, in contrast to the expectations, rhizosheath mass was positively impacted, when a large proportion of roots grew into macropores (Fig. 7D). These results probably reflect soil being glued together by plant-released organic compounds and root hairs (York *et al*., 2022; Aslam *et al*., 2022). Although root hairs were not measured here, previous studies suggest that they develop in patchy patterns, and can be particularly long in areas with low soil contact, such as macropores (Duddek *et al*., 2024). The increase in plant-originated carbon found within the rhizosheath (Fig. 7C) might in part reflect the presence of detached root hairs when the plant grew into the soil macropores (Fig. 7D). It is important to note that our imaging resolution did not permit visualization of root hairs or their contribution to root-soil contact.

In line with these findings, our third hypothesis, was not supported, as we found no relationship (*P. virgatum*) or even partly negative (*R. hirta*) relationship between rhizosheath mass and the X-ray CT derived rhizosphere density (Fig. 7B). This lack of association might be primarily the result of rhizosheath formation being associated with roots growing into macropores (Fig. 7D), while the most dense rhizosphere was found for roots which have grown into the soil matrix (Fig. S. S3A, B). Thus, rhizosheath mass and X-ray CT derived rhizosphere density for the roots of *R. hirta* were even negatively correlated (Fig. 7B). Similarly, Rabbi et al. (2018b) observed increased macroporosity near roots and higher rhizosheath mass in drought-tolerant species compared to drought-sensitive species, attributing these differences to mucilage acting as a glue between soil particles and stabilize particles during drying.

In summary, and contrary to the common assumption, our data does not support the notion that rhizosheath mass is a proxy for rhizosphere structure or soil contact in general. Rather, the rhizosheath seems to form mainly around roots that grow in macropores and thus possibly have more root hairs to improve soil contact. In contrast to roots growing in macropores, which have reduced soil contact, roots penetrating the soil matrix maintain a high degree of root–soil contact (Wendel *et al*., 2022; Lucas *et al*., 2023b). Thus, roots growing into the soil matrix might not depend on root-hairs, and in such cases the rhizosphere tends to be poorly attached to the roots and is easily detached during rhizosheath sampling. While the rhizosheath mass may indicate improved soil contact in high-macroporosity soils, such as sands (Price, 1911; Buckley, 1982), our results suggest it is not representative for the rhizosphere of an entire root system in finer-structured soils.

### 4.4 Implications for further rhizosphere and rhizosheath focused research

Our study demonstrated that root interactions with the inherent pore structure are pivotal in shaping rhizosphere properties, with root growth patterns − rather than direct root-induced changes − emerging as the primary driver of local macroporosity gradients in intact soils. Unlike the homogenized structure of sieved soils commonly used in experimental settings, intact field soils typically contain a complex network of natural macropores. In other words, in sieved soils plant roots have to heavily rearrange the existing structure and induce changes to the local macroporosity, while in intact soil these changes were conducted by previous plants or management practices. Hence, root-induced changes often reported in experiments with sieved soils are likely overestimated when compared to what takes place in the field. This highlights the necessity of conducting studies in intact soil systems to better capture the complexities of rhizosphere formation under natural conditions.

Due to challenges in rhizosphere sampling and quantification, researchers often use rhizosheath soil as a rhizosphere proxy and compare its properties to the bulk soil (Hallett *et al*., 2022; Mueller *et al*., 2024; Steiner *et al*., 2024, 2025). This approach assumes that root-adhering soil is representative of the entire rhizosphere (Pang *et al*., 2017), yet our findings challenge the validity of this assumption. Future research should integrate structural analyses of the rhizosphere with rhizosheath measurements to find optimal strategies for holistic assessments of physical, chemical, and biological properties of the entire rhizosphere.

## 5 Data availability

All raw image data that support this study are available under https://doi.org/10.48758/ufz.13657.

## 6 Acknowledgements

This research was funded in part by the Great Lakes Bioenergy Research Center, U.S. Department of Energy, Office of Science, Office of Biological and Environmental Research under Award Number DE-SC0018409, by the NSF DEB Program (Award # 1904267), by the NSF LTER Program (DEB 1027253) at the Kellogg Biological Station, and by Michigan State University AgBioResearch.

We thank Maxwell Oerther for his great support during the whole experiment, Jinho Lee for his help on the field site and Michelle Quigley for her support during the X-ray µCT scans. We highly appreciate the help from the laboratories of Eric Patterson and Thomas Sharkey during the 14C analysis and 14C labelling, respectively.

## 8 Author contributions

A.G. and M.L. constructed and built the plant pots, ingrowth cores and cores for collecting soil water. M.L., A.K. and A.G. designed the experimental setup. M.L conducted the experimental work. All authors discussed the results and finalized the paper. A.K. and M.L. analyzed the data and wrote the paper.

## References

Aravena JE, Berli M, Ghezzehei TA, Tyler SW. 2011. Effects of root-induced compaction on rhizosphere hydraulic properties--X-ray microtomography imaging and numerical simulations. Environmental science & technology 45: 425–31.

Aslam MM, Karanja JK, Dodd IC, Waseem M, Weifeng X. 2022. Rhizosheath: An adaptive root trait to improve plant tolerance to phosphorus and water deficits? Plant, Cell & Environment 45: 2861–2874.

Bodner G, Leitner D, Kaul H-P. 2014. Coarse and fine root plants affect pore size distributions differently. Plant and soil 380: 133–151.

Bruand A, Cousin I, Niccoullaud B, Duval O, Begon JC. 1996. Backscattered Electron Scanning Images of Soil Porosity for Analyzing Soil Compaction around Roots. Soil Science Society of America Journal 60: 895–901.

Buckley R. 1982. Sand rhizosheath of an arid zone grass. Plant and Soil 66: 417–421.

Carminati A, Vetterlein D, Koebernick N, Blaser S, Weller U, Vogel H-J. 2013. Do roots mind the gap? Plant and Soil 367: 651–661.

Cheraghi M, Mousavi SM, Zarebanadkouki M. 2023. Functions of rhizosheath on facilitating the uptake of water and nutrients under drought stress: A review. Plant and Soil.

Dexter AR. 1987. Compression of soil around roots. Plant and Soil 97: 401–406.

Duddek P, Papritz A, Ahmed MA, Lovric G, Carminati A. 2024. Observations of root hair patterning in soils: Insights from synchrotron-based X-ray computed microtomography. Plant and Soil.

Feeney DS, Crawford JW, Daniell T, Hallett PD, Nunan N, Ritz K, Rivers M, Young IM. 2006. Three-dimensional microorganization of the soil-root-microbe system. Microbial ecology 52: 151–8.

Goerg GM. 2015. The Lambert Way to Gaussianize Heavy-Tailed Data with the Inverse of Tukey’s *h* Transformation as a Special Case (T Hu, Ed.). The Scientific World Journal 2015: 909231.

Hallett PD, Marin M, Bending GD, George TS, Collins CD, Otten W. 2022. Building soil sustainability from root–soil interface traits. Trends in Plant Science 27: 688–698.

Helliwell JR, Sturrock CJ, Mairhofer S, Craigon J, Ashton RW, Miller AJ, Whalley WR, Mooney SJ. 2017. The emergent rhizosphere: imaging the development of the porous architecture at the root-soil interface. Scientific reports 7: 14875.

Helliwell JR, Sturrock CJ, Miller AJ, Whalley WR, Mooney SJ. 2019. The role of plant species and soil condition in the structural development of the rhizosphere. Plant, cell & environment 42: 1974–1986.

Hildebrand T, Rüegsegger P. 1997. A new method for the model-independent assessment of thickness in three-dimensional images. Journal of Microscopy 185: 67–75.

Hinsinger P, Bengough AG, Vetterlein D, Young IM. 2009. Rhizosphere: biophysics, biogeochemistry and ecological relevance. Plant and Soil 321: 117–152.

Koebernick N, Daly KR, Keyes SD, Bengough AG, Brown LK, Cooper LJ, George TS, Hallett PD, Naveed M, Raffan A, et al. 2019. Imaging microstructure of the barley rhizosphere: particle packing and root hair influences. The New phytologist 221: 1878–1889.

Koebernick N, Daly KR, Keyes SD, George TS, Brown LK, Raffan A, Cooper LJ, Naveed M, Bengough AG, Sinclair I, et al. 2017. High-resolution synchrotron imaging shows that root hairs influence rhizosphere soil structure formation. The New phytologist 216: 124–135.

Koebernick N, Schlüter S, Blaser SRGA, Vetterlein D. 2018. Root-soil contact dynamics of Vicia faba in sand. Plant and Soil 431: 417–431.

Kravchenko AN, Guber AK, Razavi BS, Koestel J, Quigley MY, Robertson GP, Kuzyakov Y. 2019. Microbial spatial footprint as a driver of soil carbon stabilization. Nature communications 10: 3121.

Kutilek M, Jendele L, Panayiotopoulos K. 2006. The influence of uniaxial compression upon pore size distribution in bi-modal soils. Soil and Tillage Research 86: 27–37.

Kuzyakov Y, Razavi BS. 2019. Rhizosphere size and shape: Temporal dynamics and spatial stationarity. Soil Biology and Biochemistry 135: 343–360.

Landl M, Phalempin M, Schlüter S, Vetterlein D, Vanderborght J, Kroener E, Schnepf A. 2021. Modeling the Impact of Rhizosphere Bulk Density and Mucilage Gradients on Root Water Uptake. Frontiers in Agronomy 3.

Lenth RV. 2024. emmeans: Estimated Marginal Means, aka Least-Squares Means.

Li Z, Kravchenko AN, Cupples A, Guber AK, Kuzyakov Y, Philip Robertson G, Blagodatskaya E. 2024. Composition and metabolism of microbial communities in soil pores. Nature Communications 15: 3578.

Lucas M. 2020. Soil structure formation through the action of plants. Universitäts- und Landesbibliothek Sachsen-Anhalt.

Lucas M, Gil J, Robertson GP, Ostrom NE, Kravchenko A. 2023a. Changes in soil pore structure generated by the root systems of maize, sorghum and switchgrass affect in situ N2O emissions and bacterial denitrification. Biology and Fertility of Soils.

Lucas M, Nguyen LTT, Guber A, Kravchenko AN. 2022. Cover crop influence on pore size distribution and biopore dynamics: Enumerating root and soil faunal effects. Frontiers in plant science 13: 928569.

Lucas M, Santiago JP, Chen J, Guber A, Kravchenko A. 2023b. The soil pore structure encountered by roots affects plant-derived carbon inputs and fate. New Phytologist 240: 515–528.

Lucas M, Schlüter S, Vogel H-J, Vetterlein D. 2019. Roots compact the surrounding soil depending on the structures they encounter. Scientific reports 9: 16236.

Marschner P. 2012. Rhizosphere Biology. In: Marschner’s Mineral Nutrition of Higher Plants. Elsevier, 369–388.

Mo X, Wang M, Zeng H, Wang J. 2023. Rhizosheath: Distinct features and environmental functions. Geoderma 435: 116500.

Mueller CW, Baumert V, Carminati A, Germon A, Holz M, Kögel-Knabner I, Peth S, Schlüter S, Uteau D, Vetterlein D, et al. 2024. From rhizosphere to detritusphere – Soil structure formation driven by plant roots and the interactions with soil biota. Soil Biology and Biochemistry: 109396.

Ndour PMS, Heulin T, Achouak W, Laplaze L, Cournac L. 2020. The rhizosheath: from desert plants adaptation to crop breeding. Plant and Soil 456: 1–13.

Pang J, Ryan MH, Siddique KHM, Simpson RJ. 2017. Unwrapping the rhizosheath. Plant and Soil 418: 129–139.

Pankievicz VCS, Delaux P-M, Infante V, Hirsch HH, Rajasekar S, Zamora P, Jayaraman D, Calderon CI, Bennett A, Ané J-M. 2022. Nitrogen fixation and mucilage production on maize aerial roots is controlled by aerial root development and border cell functions. Frontiers in Plant Science 13: 977056.

Pausch J, Kuzyakov Y. 2011. Photoassimilate allocation and dynamics of hotspots in roots visualized by 14 C phosphor imaging. Journal of Plant Nutrition and Soil Science 174: 12–19.

Phalempin M, Lippold E, Vetterlein D, Schlüter S. 2021. Soil texture and structure heterogeneity predominantly governs bulk density gradients around roots. Vadose Zone Journal 20.

Phalempin M, Rosskopf U, Schlüter S, Vetterlein D, Peth S. 2023. Can we use X-ray CT to generate 3D penetration resistance data? Geoderma 439: 116700.

Pinheiro JC, Bates DM. 2000. Mixed-Effects Models in S and S-PLUS. New York: Springer.

Pinheiro J, Bates D, R Core Team. 2024. nlme: Linear and Nonlinear Mixed Effects Models.

Price SR. 1911. THE ROOTS OF SOME NORTH APRICAN DESERT-GRASSES. New Phytologist 10: 328–340.

R Core Team. 2023. R: A Language and Environment for Statistical Computing. Vienna, Austria: R Foundation for Statistical Computing.

Rabbi SMF, Tighe MK, Flavel RJ, Kaiser BN, Guppy CN, Zhang X, Young IM. 2018a. Plant roots redesign the rhizosphere to alter the three-dimensional physical architecture and water dynamics. The New phytologist 219: 542–550.

Rabbi SMF, Tighe MK, Knox O, Young IM. 2018b. The impact of carbon addition on the organisation of rhizosheath of chickpea. Scientific reports 8: 18028.

Ruamps LS, Nunan N, Pouteau V, Leloup J, Raynaud X, Roy V, Chenu C. 2013. Regulation of soil organic C mineralisation at the pore scale. FEMS Microbiology Ecology 86: 2635.

Rueden CT, Schindelin J, Hiner MC, DeZonia BE, Walter AE, Arena ET, Eliceiri KW. 2017. ImageJ2: ImageJ for the next generation of scientific image data. BMC bioinformatics 18: 529.

Santiago JP, Soltani A, Bresson MM, Preiser AL, Lowry DB, Sharkey TD. 2021. Contrasting anther glucose-6-phosphate dehydrogenase activities between two bean varieties suggest an important role in reproductive heat tolerance. Plant, cell & environment 44: 2185–2199.

Schindelin J, Arganda-Carreras I, Frise E, Kaynig V, Longair M, Pietzsch T, Preibisch S, Rueden C, Saalfeld S, Schmid B, et al. 2012. Fiji: an open-source platform for biological-image analysis. Nature Methods 9: 676–682.

Schmidt H, Nunan N, Höck A, Eickhorst T, Kaiser C, Woebken D, Raynaud X. 2018. Recognizing Patterns: Spatial Analysis of Observed Microbial Colonization on Root Surfaces. Frontiers in Environmental Science 6: 61.

Schnepf A, Carminati A, Ahmed MA, Ani M, Benard P, Bentz J, Bonkowski M, Knott M, Diehl D, Duddek P, et al. 2022. Linking rhizosphere processes across scales: Opinion. Plant and Soil 478: 5–42.

Steiner FA, Tung S-Y, Wild AJ, Köhler T, Tyborski N, Carminati A, Pausch J, Lüders T, Wolfrum S, Mueller CW, et al. 2025. Soil drying shapes rhizosheath properties and their link with maize yields across different soils. Plant and Soil.

Steiner FA, Wild AJ, Tyborski N, Tung S, Koehler T, Buegger F, Carminati A, Eder B, Groth J, Hesse BD, et al. 2024. Rhizosheath drought responsiveness is variety-specific and a key component of belowground plant adaptation. New Phytologist 242: 479–492.

van Veelen A, Koebernick N, Scotson CS, McKay-Fletcher D, Huthwelker T, Borca CN, Mosselmans JFW, Roose T. 2019. Root-induced soil deformation influences Fe, S and P: rhizosphere chemistry investigated using synchrotron XRF and XANES. New Phytologist.

Vogel H-J, Weller U, Schlüter S. 2010. Quantification of soil structure based on Minkowski functions. Computers & Geosciences 36: 12361245.

Vogel H-J, Weller U, Schlüter S. 2024. Linking structure and functions in agricultural soils. In: Advances in Agronomy. Elsevier, 363–403.

Vollsnes AV, Futsaether CM, Bengough AG. 2010. Quantifying rhizosphere particle movement around mutant maize roots using time-lapse imaging and particle image velocimetry. European Journal of Soil Science 61: 926–939.

Watt M, McCully ME, Canny MJ. 1994. Formation and Stabilization of Rhizosheaths of Zea mays L.’. 106.

Wendel AS, Bauke SL, Amelung W, Knief C. 2022. Root-rhizosphere-soil interactions in biopores. Plant and Soil.

White RG, Kirkegaard JA. 2010. The distribution and abundance of wheat roots in a dense, structured subsoil--implications for water uptake. Plant, cell & environment 33: 133–48.

York LM, Cumming JR, Trusiak A, Bonito G, Von Haden AC, Kalluri UC, Tiemann LK, Andeer PF, Blanc-Betes E, Diab JH, et al. 2022. Bioenergy Underground: Challenges and opportunities for phenotyping roots and the microbiome for sustainable bioenergy crop production. The Plant Phenome Journal 5: e20028.

Young IM, Crawford JW. 2004. Interactions and Self-Organization in the Soil-Microbe Complex. Science 304: 1634–1637.

Zhang Y, Du H, Gui Y, Xu F, Liu J, Zhang J, Xu W. 2020. Moderate water stress in rice induces rhizosheath formation associated with abscisic acid and auxin responses. Journal of experimental botany.

