## Supplementary material for "Root-pore interactions, the underestimated driver for rhizosphere structure and rhizosheath development": Suplementary File

The following Supporting Information is available for this article:

**Fig. S1 Examples of X-ray µCT images of roots growing into macropores, biopores, and soil matrix devoid of macropores.**

**Fig. S2 Plots of residuals versus fitted values for the model with A) changes in macropores and B) changes in narrow pores as the response variable.**

**Fig. S3 Associations of the effects on macroporosity from root-induced changes and root growth preferences with the share of roots grown into the soil matrix.**

**Fig. S4 Narrow pores (0.04-0.15 mm diameter) as a function of distance to the root surface.**

**Fig. S5 Association between root-growth preference effect at <1 mm distance from the root and changes in total macroporosity.**

**Fig. S6X-ray µCT grey values as a function of distance to the root surface.**

**Fig. S7 Association between the share of roots grown either in biopores or the soil matrix and the rhizosheath dry weight.**

**Table S1 Results of ANOVA for rhizosphere macroporosity and for narrow pores.**

**Methods S1 X-ray CT of ingrowth cores and image segmentation**

**Fig. S1 Examples of X-ray µCT images of roots growing into macropores, biopores, and soil matrix devoid of macropores.** A) Root (orange) grown into a large macropore filled with air (black), B) root grown into a cylindrical biopore and C) root grown through the dense soil matrix (grey). Note that in our analysis of local rhizosphere macroporosity, biopores are treated as a subset of all macropores.

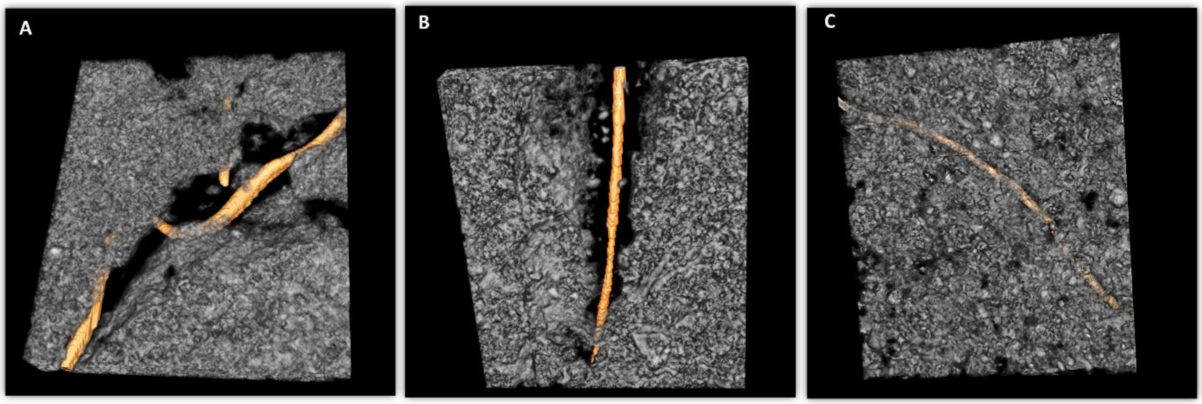


**Fig. S2 Plots of residuals versus fitted values for the model with A) changes in macropores and B) changes in narrow pores as the response variable.**

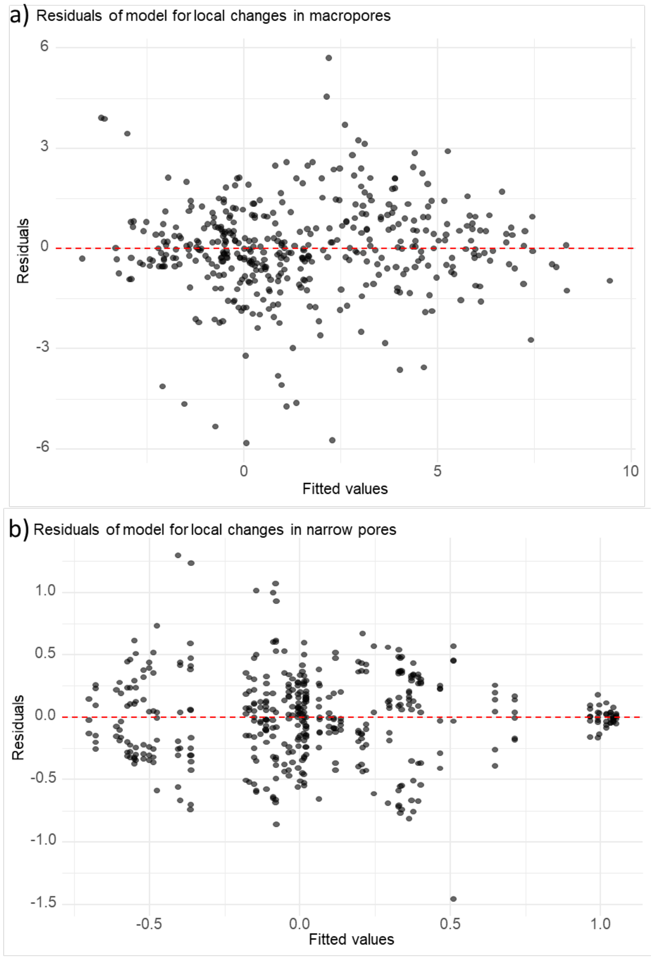

**Fig. S3 Associations of the effects on macroporosity from root-induced changes and root growth preferences with the share of roots grown into the soil matrix.** Associations of the root growth preference effect at <0.2 mm (a) and <1 mm (b) distance to the root with the share of roots grown into the soil matrix and association of root-induced changes at <0.2 mm (c) and <1 mm (d) distance to the root with the share of roots grown into the soil matrix. Solid lines indicate significant linear associations, while a dashed line indicates non-significant associations between the depicted variables (p-value>0.05).

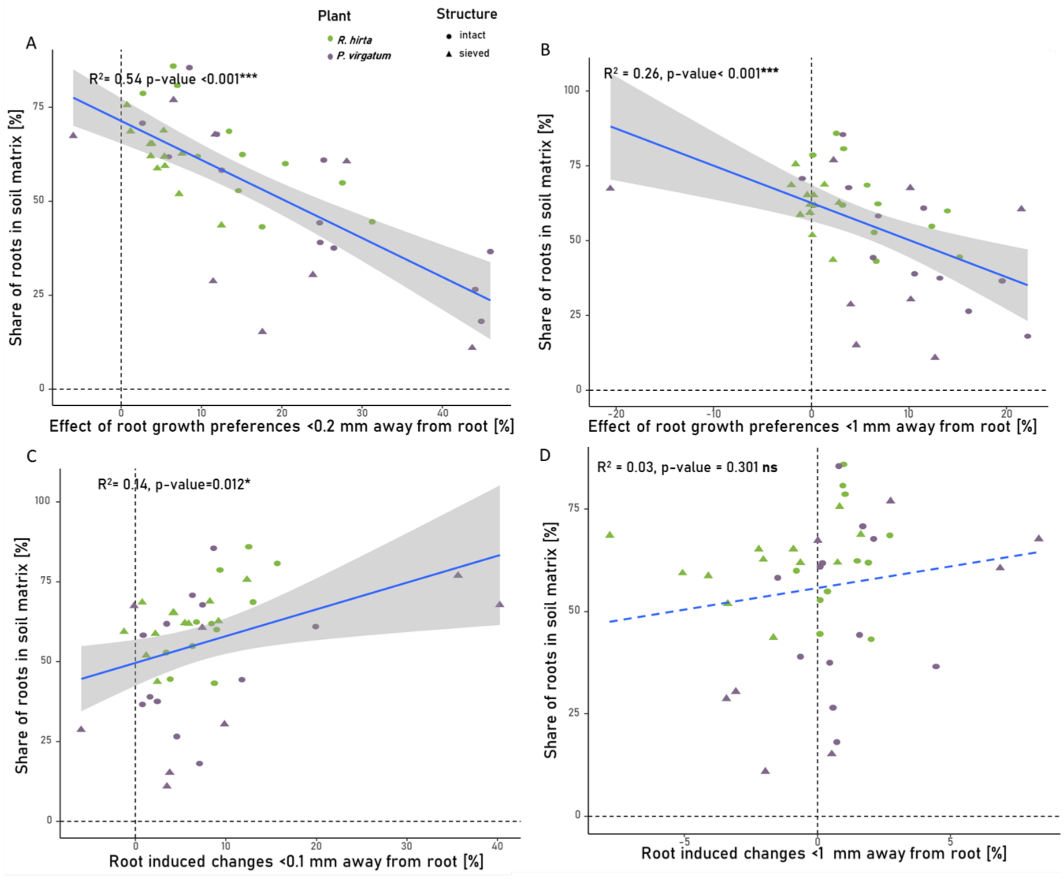

**Fig. S4 Narrow pores (0.04-0.15 mm diameter) as a function of distance to the root surface.** Gradients of narrow pores with distance to the root surface of Panicum virgatum and Rudbeckia hirta when grown into sieved or intact soil of prairie or switchgrass origin. Depicted are the local volumes of narrow pores as percentage of the total local volume computed before the plants were seeded (dotted line) and after roots developed. In addition, the mean share of narrow pores (per sample) is shown (dashed line).

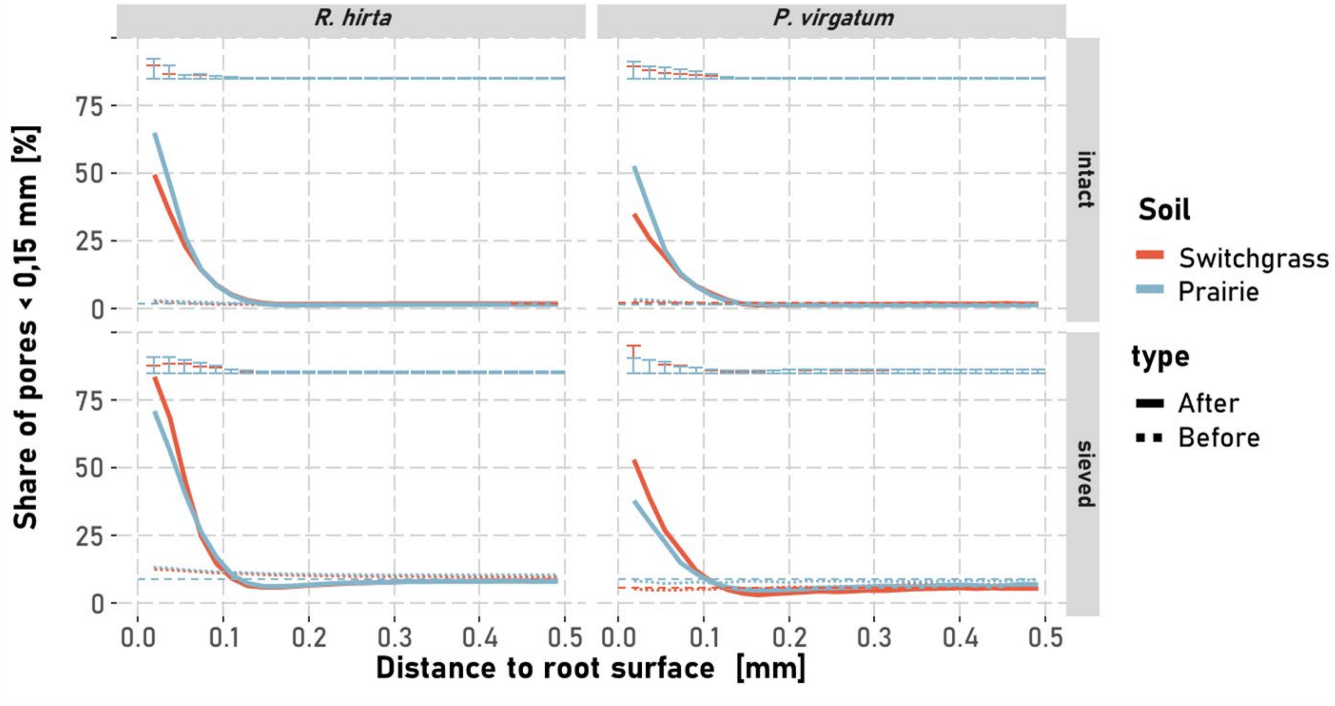

**Fig. S5 Association between root-growth preference effect at <1 mm distance from the root and changes in total macroporosity.**

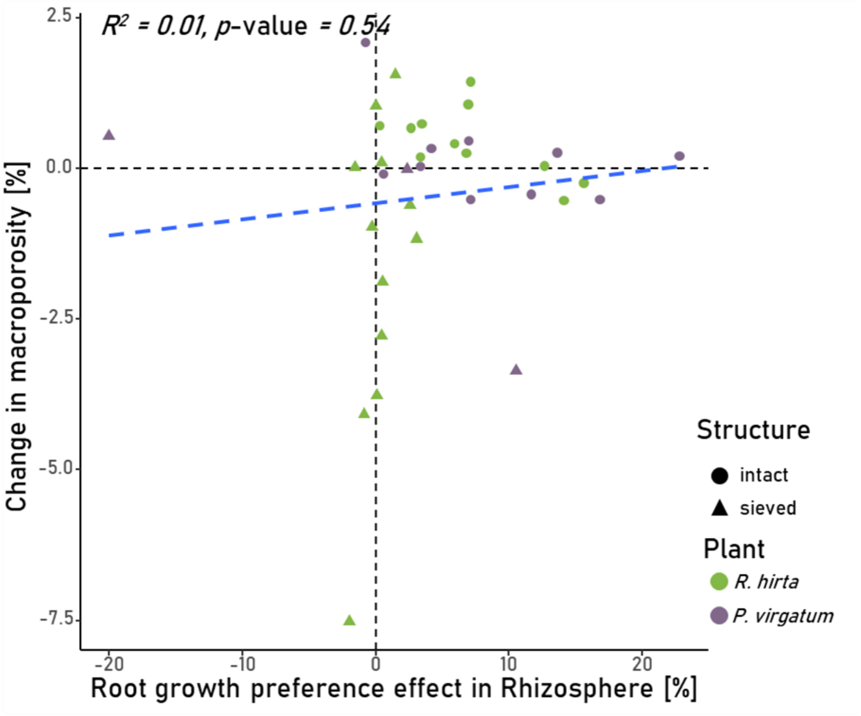

**Fig. S6 X-ray µCT grey values as a function of distance to the root surface.** Gradients of X-ray CT image grey values with distance to the root surface of Panicum virgatum and Rudbeckia hirta when grown into sieved or intact soil of prairie or switchgrass origin. Depicted are also the mean grey values per treatment (proxy for bulk density, dashed lines).

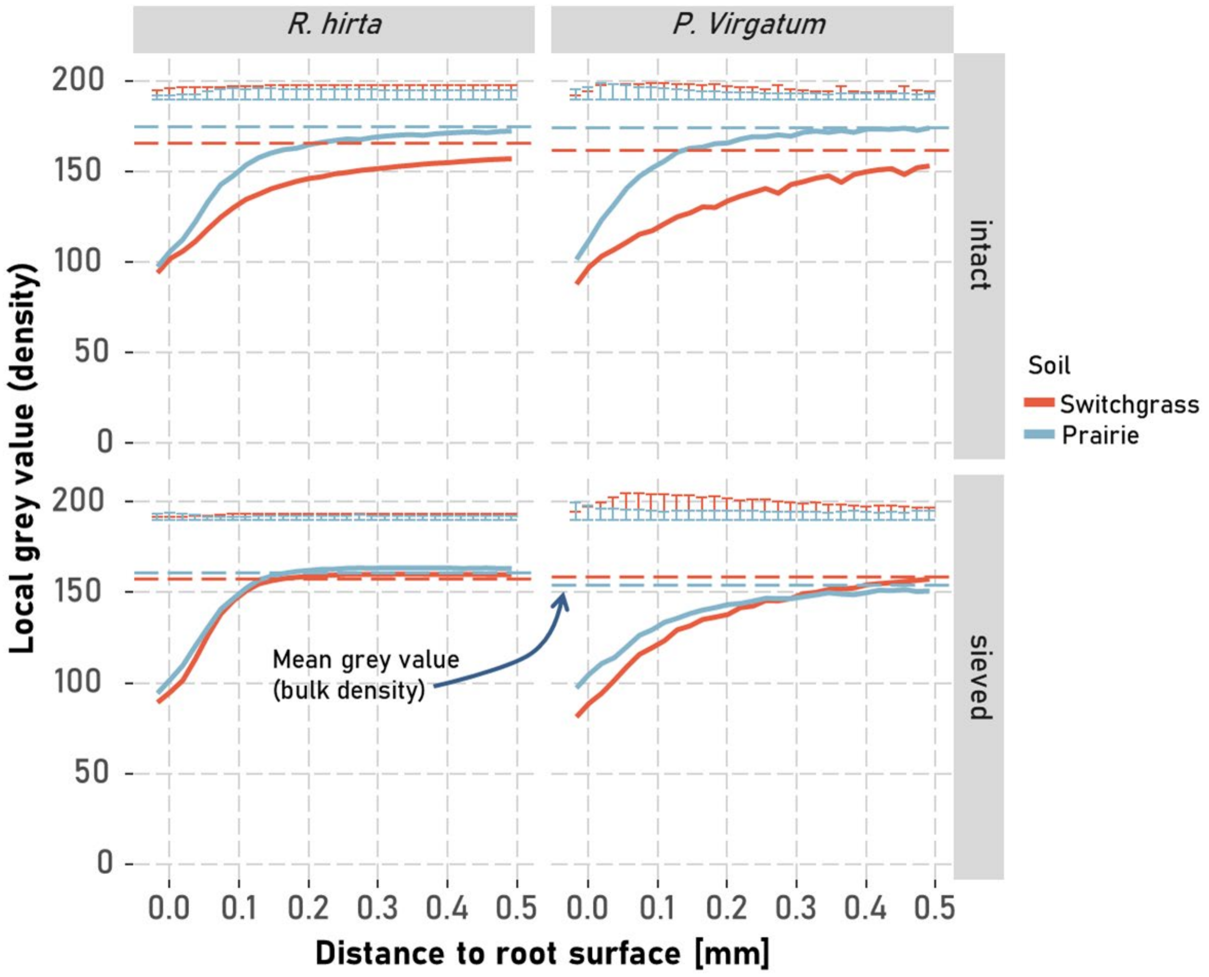

**Fig. S7 Association between the share of roots grown either in biopores or the soil matrix and the rhizosheath dry weight.** A) Association of the share of roots found in biopores with the rhizosheath dry weight and B) association of the share of roots found in the soil matrix with the rhizosheath dry weight. Solid and dashed lines represent linear regression models fitted to the data that were, respectively, statistically significant or not at p-value <0.05.

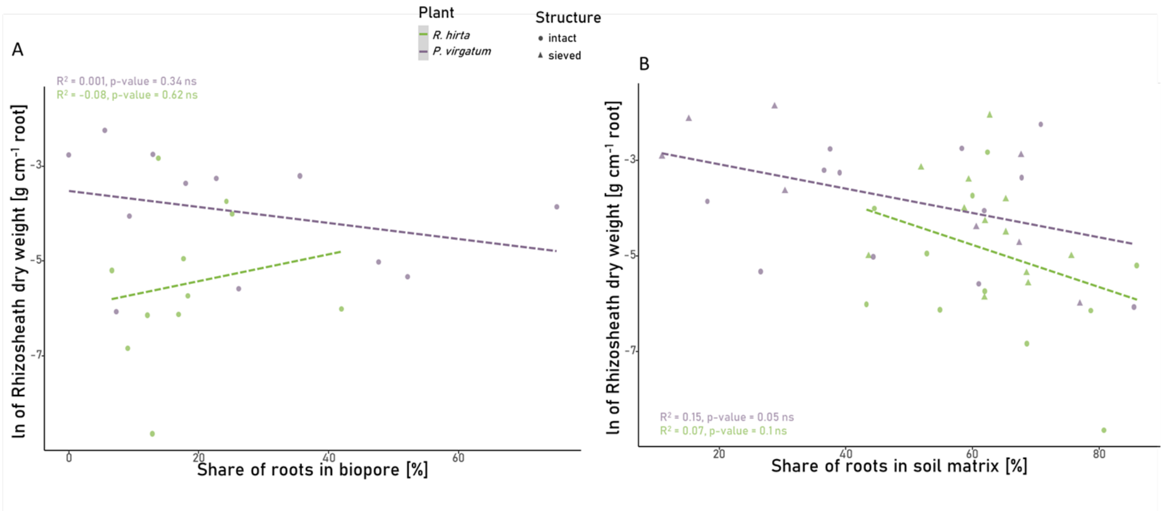

**Table S1 Results of ANOVA for rhizosphere macroporosity and for narrow pores** Shown are the F and p-values for the four studied factors, namely, plant (*P. virgatum* vs. *R. hirta*), soil structure (sieved vs. intact), soil origin (switchgrass soil vs. prairie soil), and the type of process leading to variations in local macroporosity (root-induced changes vs. root growth preferences), and their interactions. Significant effects (p-value < 0.05) are marked in bold.

| | **macropores > 0.04 mm** | | **narrow pores 0.04-0.15 mm** | |
|---|---|---|---|---|
| | F-ratio | P value | F-ratio | p-value |
| **Plant** | **4.314** | **0.055** | **11.003** | **0.005** |
| **Structure** | **17.139** | **0.009** | **11.169** | **0.021** |
| **Soil** | 0.045 | 0.837 | 0.835 | 0.382 |
| **Process** | **182.820** | **<0.001** | **5.315** | **0.022** |
| **Distance_mm** | **55.273** | **<0.001** | **52.64** | **<0.001** |
| **Plant:Structure** | 4.137 | 0.060 | **7.16** | **0.017** |
| **Plant:Soil** | 0.261 | 0.617 | 0.283 | 0.603 |
| **Plant:Process** | **13.805** | **<0.001** | 1.618 | 0.204 |
| **Plant:Distance_mm** | 1.887 | 0.113 | 1.71 | 0.148 |
| **Structure:Soil** | 5.772 | 0.037 | 2.882 | 0.120 |
| **Structure:Process** | 6.277 | 0.013 | **90.272** | **<0.001** |
| **Structure:Distance_mm** | 0.217 | 0.929 | 0.442 | 0.778 |
| **Soil:Process** | **42.696** | **<0.001** | 1.415 | 0.235 |
| **Soil:Distance_mm** | 0.074 | 0.990 | 0.028 | 0.998 |
| **Process:Distance_mm** | **6.179** | **<0.001** | **33.179** | **<0.001** |
| **Plant:Structure:Soil** | 0.858 | 0.369 | 1.183 | 0.294 |
| **Plant:Structure:Process** | **5.151** | **0.024** | **16.894** | **<0.001** |
| **Plant:Structure:Distance_mm** | 1.704 | 0.149 | 1.579 | 0.180 |
| **Plant:Soil:Process** | 5.391 | 0.021 | 3.127 | 0.078 |
| **Plant:Soil:Distance_mm** | 0.174 | 0.952 | 0.159 | 0.959 |
| **Plant:Process:Distance_mm** | 0.393 | 0.814 | 0.481 | 0.750 |
| **Structure:Soil:Process** | **13.682** | **<0.001** | **5.407** | **0.021** |
| **Structure:Soil:Distance_mm** | 0.406 | 0.804 | 0.759 | 0.552 |
| **Structure:Process:Distance_mm** | 0.329 | 0.858 | **4.164** | **0.003** |
| **Soil:Process:Distance_mm** | 0.694 | 0.597 | 0.836 | 0.503 |
| **Plant:Structure:Soil:Process** | 3.324 | 0.069 | 1.211 | 0.272 |
| **Plant:Structure:Soil:Distance_mm** | 0.205 | 0.936 | 0.342 | 0.849 |
| **Plant:Structure:Process:Distance_mm** | 0.882 | 0.475 | 0.038 | 0.997 |
| **Plant:Soil:Process:Distance_mm** | 0.276 | 0.893 | 0.55 | 0.699 |
| **Structure:Soil:Process:Distance_mm** | 0.073 | 0.990 | 0.1 | 0.982 |
| **Plant:Structure:Soil:Process:Distance_mm** | 0.163 | 0.957 | 0.152 | 0.962 |

Model structure: lme(Macropore Change~ Plant*Structure*Soil*Process*Distance_mm, random = ~1|Plant_replicate/Structure/Soil/Core, weights= varComb(varIdent(form = ~1|Plant), varIdent(form = ~1|Distance_mm), model_data)

### Methods S1 X-ray CT of ingrowth cores and image segmentation

All ingrowth cores were scanned using an X-ray microtomograph (X3000, North Star Imaging, USA) at 75 kV and 470 µA, both before and after the growth experiment. Despite the energy settings causing a larger focal spot on the Varian L07 detector (1920x1536 pixels), a SubpiX mode provided a voxel size of 18.2 µm. Each scan consisted of four sub-images (2 rows, 2 columns), capturing 2880 projections at 3 fps, averaging 2 frames. 3D image reconstruction was performed using efX software. To track root growth, X-ray CT images taken before and after the experiment were registered using Elastix software (Klein *et al.*, 2010; Shamonin, 2013) as described by Lucas et al. (2020). The registered soil images were cropped to 1850x1850x2300 voxels in Fiji (Ollion *et al.*, 2013) to eliminate artifacts along the core walls. Contrast was enhanced (saturation = 0.35), bit depth reduced to 8-bit, and a non-local means filter (Darbon *et al.*, 2008; Buades *et al.*, 2011) was applied using scikit-image (van der Walt et al., 2014) in Python (van Rossum & Drake, 2009). Macropores (>40 µm) were separated from the soil matrix using the Otsu algorithm (Otsu, 1979). Biopores and roots were segmented according to Lucas et al. (2022), with scripts available on GitHub ('https://github.com/Maik-Lu/Roots_and_Biopores'). In short, the scripts rely on the Tubeness plugin in Fiji to separate different sized tubular objects from the remaining irregularly shaped pore network. New roots were identified by subtracting the roots in the pre-experiment images from those in the post-experiment images.
